# Cryptic Variation in Adaptive Phenotypes Revealed by Panspecific *flc* Mutants

**DOI:** 10.1101/2024.06.14.599000

**Authors:** Ulrich Lutz, Ilja Bezrukov, Rebecca Schwab, Wei Yuan, Marius Kollmar, Detlef Weigel

**Author notes:** Department of Physiology of Yield Stability, Faculty of Agriculture, University of Hohenheim, Stuttgart, Germany (Marius Kollmar).

## Abstract

The study of mutants is one of the best tools for understanding the genetic basis of phenotypes that contribute to adaptation. Oddly, mutant analyses are almost always restricted to single genetic backgrounds and findings therefore can not be easily generalized. A case in point is the key regulator of flowering, *FLOWERING LOCUS C* (*FLC*), which has been inferred to explain much of the flowering time variation in *Arabidopsis thaliana*, yet mutants have been examined in very few backgrounds. We have previously established a set of panspecific *flc* mutants in 62 accessions of *A. thaliana* (Ruffley et al. 2024). Here, we investigate how genetic background modulates mutant effects on flowering and vegetative traits, as well as on physiology and transcriptomes. Time to onset of flowering in the genome-edited *flc* lines was reduced by up to 83%, but considerable variation remained. Genetic mapping showed that extremely early flowering in the absence of *FLC* was mostly explained by natural variation at the known FLC target *FT*, with additional contribution from loci colocalizing with *FLC*. Prognostic sequence analyses of accessions did not suggest that extremely-early combinations of engineered *flc* and natural *FT* alleles would be deleterious, yet extremely early flowering accessions are not represented in the commonly used collections of *A. thaliana* accessions. To test whether this discrepancy could be due to sampling bias, we undertook a focused collection effort of wild populations in Southern Italy, which confirmed that extremely early flowering accessions exist in natural populations. Apart from its specific role in flowering time regulation, *FLC* has pleiotropic effects on other ecophysiological traits such as growth, and these were also dependent on the genetic background, which was further supported by transcriptomic comparisons. Together we conclude that the various roles of *FLC* have greatly diversified in different genetic backgrounds. Our study provides a proof-of-concept on how analysis of panspecific mutants can reveal the true extent of genetic networks in which a focal gene participates in.

## Introduction

The onset of flowering is an important developmental transition resulting from the coupling of an internal genetic program to external local and seasonal climate cues. Its correct timing is highly adaptive and greatly affects fitness. In *Arabidopsis thaliana*, its molecular control is orchestrated by a large network of genes integrating a multitude of signals that converge on a few key regulators (Blümel et al. 2015; Cho et al. 2017a). *FLOWERING LOCUS C* (*FLC*), encoding a MADS-box transcription factor, is a central floral repressor that prevents flowering before favorable spring conditions to integrate with a cold winter season by vernalization. Among many direct transcriptional targets, it acts through the repression of floral inducers like *FLOWERING LOCUS T* (*FT*) and *SUPPRESSOR OF CONSTANS 1* (*SOC1*) (Michaels and Amasino 1999; Amasino 2010; Deng et al. 2011). In winter-annual accessions of *A. thaliana*, flowering is prevented by high *FLC* levels until a long period of winter cold, or vernalization, caused by its activation through the FRIGIDA (FRI) supercomplex, that recruits transcription factors and chromatin modifiers (Lee et al. 1993; Sheldon et al. 2000; Shindo et al. 2005; Hepworth et al. 2018, 2020). Before vernalization, the chromatin of *FLC* is highly enriched with activating histone marks. Gradual chromatin deacetylation and removal of histone trimethylation (H3K27me3) during sustained cold temperature, together with a decoration of the *FLC* nucleation region with repressive histone marks lead to an overall and stable suppression of *FLC* upon the increase of ambient growth temperature (Finnegan and Dennis 2007; Lucia et al. 2008; Angel et al. 2011; Coustham et al. 2012; Li et al. 2014; Yang et al. 2017).

Extensive natural variation in vernalization sensitivity and flowering time is present in accessions of *A. thaliana* and many quantitative trait loci (QTL) have been identified that contribute to this variation. Among these, QTLs that map to *FRI* or *FLC* explain a large fraction of the variation, even after vernalization treatments, probably due to these treatments often not being saturating. The strong effects of *FRI* and *FLC* in many cases mask the effects of other loci whilst additivity was the predominant mode of action of these other QTLs (Alonso-Blanco et al. 1998; O’Neill et al. 2008; Simon et al. 2008; Schwartz et al. 2009; Li et al. 2010; Salomé et al. 2011; Strange et al. 2011; Grillo et al. 2013; Huang et al. 2013; Dittmar et al. 2014; Sasaki et al. 2015). Natural loss-of-function allelic variants of *FRI* are widespread and positively selected, whereas complete inactivation of *FLC* is rare. Instead transposon insertions into the first intron that attenuate *FLC* expression are common (Werner et al. 2005; Schmalenbach et al. 2014; Zhang and Jiménez-Gómez 2020; Baduel et al. 2021). In *FLC*, most polymorphisms in the non-coding regions are associated with different vernalization requirements in terms of duration and temperature (Li et al. 2014; Zhu et al. 2023). Smaller-effect QTL at *FLC* have been revealed upon vernalization or in summer-annual accessions, but tight genetic linkage and massive allelic heterogeneity complicated the separation of the effects of *FLC* from the adjacent loci (El-Lithy 2005; Simon et al. 2008; Li et al. 2010; Salomé et al. 2011; Strange et al. 2011; Huang et al. 2013; Sasaki et al. 2015).

Pleiotropy describes the phenomenon that a single gene or an allele of a gene can control several traits (Paaby and Rockman 2013). Therefore, the evolution of adaptive phenotypes can be constrained due to antagonistic selection (positive impact on some traits but negative effects on others) on correlated traits, or the essential maintenance of combinations of functional traits. Such constraints may be expressed at different levels as pleiotropic genes may evolve divergent functions in different pathways. Reverse and quantitative genetics approaches in different organisms indicate that pleiotropy may be rarer than often assumed (Wagner et al. 2008; Wang et al. 2010). Furthermore, the classification and quantification of pleiotropically controlled traits is difficult. When mapping quantitative traits, limited recombination can make it difficult to distinguish true pleiotropy from genetic linkage or from indirect effects through trait correlations. Flowering time genes, which belong to the best studied group of genes in *A. thaliana* to date, have often emerged as having pleiotropic roles throughout different developmental stages (Auge et al. 2019). *FLC* is the best example of a flowering gene with pleiotropic roles (Auge et al. 2019). Besides flowering, FRI-dependent *FLC* activity affects germination, vegetative development, circadian rhythmicity, and drought tolerance (dehydration avoidance) (McKay et al. 2003; Edwards et al. 2006; Chiang et al. 2009; Mentzer et al. 2010; Willmann and Poethig 2011; Auge et al. 2017). Direct binding of FLC protein to promoters of hundreds of genes is consistent with broad pleiotropic effects, especially during cold stress (Deng et al. 2011; Mateos et al. 2015, 2017). In light of these many adaptive traits controlled by *FLC* it has been proposed that the view of *FLC*’s specific and primary function in flowering time needs to be revised and that *FLC* needs to be appreciated as a much broader contributor to the execution of diverse ecological strategies (Takou et al. 2019). To decipher this broader pleiotropic role and the cryptic spectrum of *FLC*-independent variation in flowering time on a species-wide level we have combined quantitative genetics with phenotypic, physiological, and transcriptomic studies. We focused on experiments in controlled conditions to showcase the enormous potential of panspecific genetic disruptions to gather a holistic view of gene function, and we provide an outlook of the possibilities offered by reverse ecological approaches to validate inferences from the greenhouse with observations in natural populations.

## Results

### Cryptic variation in flowering time revealed in *flc* mutants

#### Effects of *FLC* mutations on different flowering-related traits

We previously established a collection of panspecific *flc* mutants, including both knock out (KO) and knock down (KD) alleles (Ruffley et al. 2024), that now allowed us to investigate how variable the effects of a *FLC* mutation on flowering traits are between many genetic backgrounds. To investigate the genetic background-dependent effects of the mutations on flowering time we measured time until bolting (days to flowering, DTF) and the number of leaves initiated during the vegetative phase (rosette leaf number, RLN) under inductive long-day photoperiods (LD) without prior vernalization. We also determined the number of cauline leaves (CLN) as a proxy for the vegetative portion of the primary inflorescence, which produces branches and dictates inflorescence architecture critical for optimal seed set and fitness (Ratcliffe et al. 1999; Pouteau and Albertini 2009; Yamaguchi et al. 2014).

The average flowering time (mean [±sd]) of all *flc* mutant lines were 24.2 [±6.0] DTF, 17.8 [±8.1] RLN and 6.0 [±4.1] CLN. The mutants bolted 40.3 days earlier and generated 34.4 fewer RLN and 4.6 fewer CLN than the corresponding wild types (Fig. 1A). *FLC* transcript levels correlated moderately with all flowering traits in the mutants (Pearson’s *r*, DTF: 0.27, RLN: 0.23, CLN: 0.22, p < 0.05) but only with RLN in the wild types (*r* = 0.43, *p* < 0.01) (Fig. 1B, E, Fig. S2, and Table S1). The reference accession Col-0 grouped with the earlier fraction of the mutants with 19.3 [±0.9] DTF, 12.6 [±1.2] RLN and 3.2 [±0.6] CLN. Even the *flc* KO lines varied greatly in flowering time (range DTF: 24.1, RLN: 26.7) (Fig. 1B). *flc* lines derived from the three genomic backgrounds 7186 (Kn-0), 9908 (ESP-1-11), and 9925 (RUM-20) flowered extremely early (DTF < 16, RLN < 8, Fig. 1A, B, and Fig. S1). The Kn-0 wild type had been included as representing a summer-annual accession. Kn-0 flowered with 20.25 DTF as early as Col-0 (19.26 DTF, two sided Student’s t test, p > 0.05) despite higher *FLC* levels (mean relative expression level 20.1 a.u., Col-0: 0.97 a.u.), indicating the presence of additional activators of flowering in the Kn-0 genetic background.

**Figure 1:**
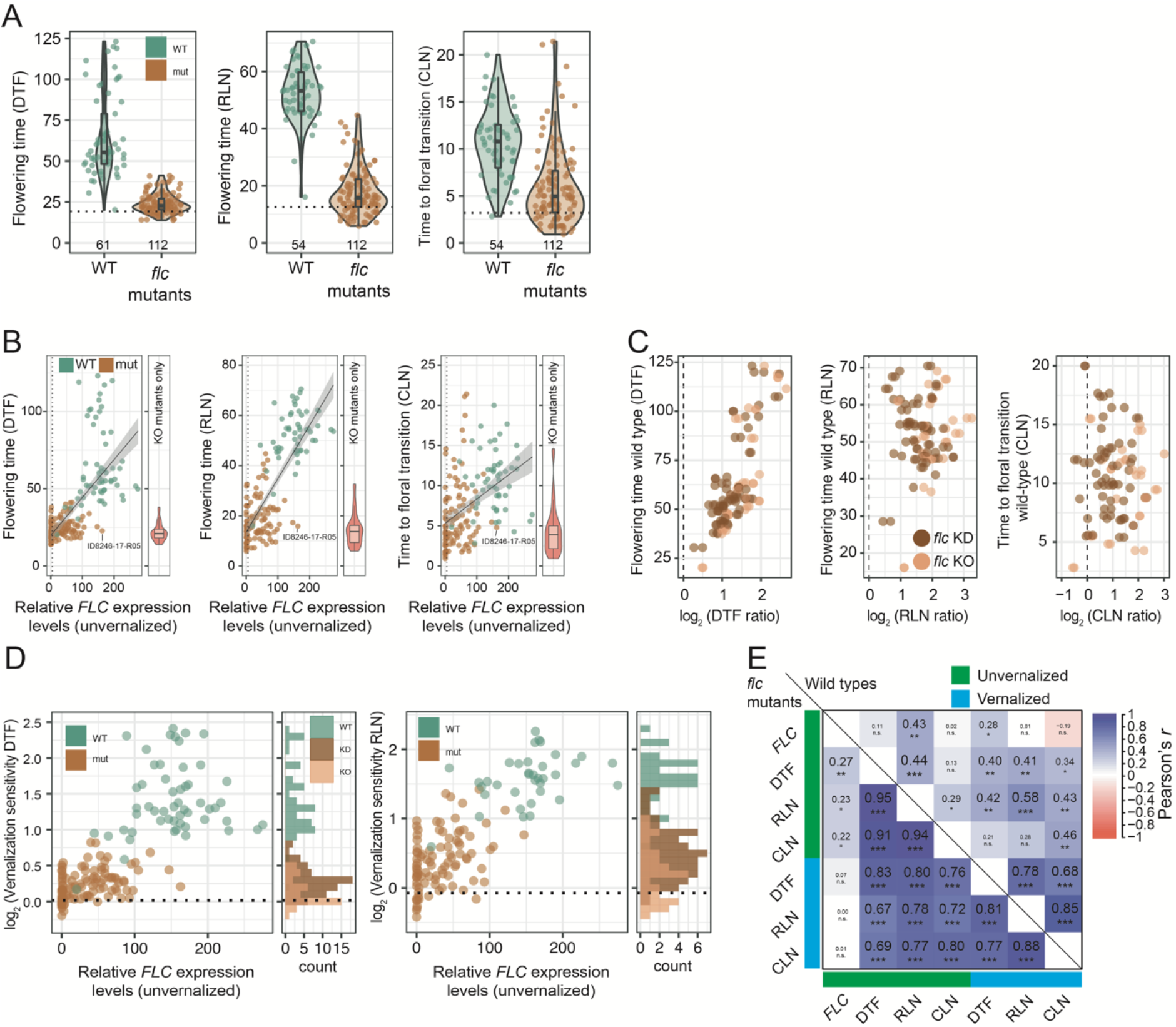
Flowering time analysis of *flc* mutants. **A**. Flowering-related traits of wild types and mutants were measured at 22°C under LD (unvernalized). Means from 3 to 12 replicates per line are shown, with the number of lines with mean values indicated at the bottom of each graph. The black dashed horizontal line indicates Col-0. **B**. Correlation between flowering time (unvernalized) and *FLC* expression levels (unvernalized). Black dashed vertical line indicates an *FLC* relative expression level of 5 a.u., which divided the mutants into KO and KD. To better illustrate the phenotypes of the KO mutants only, their values are shown as a single violin plot next to the main plot. *flc* KO, DTF mean [±sd] 21.8 [±5.6], range 13.9 to 38.0, RLN 14.6 [±6.5], range 5.9 to 32.6. **C**. Correlation of the mutant-vs.-wild type phenotypic ratios (log2[DTFWildtype / DTFMutant]) versus the wild type phenotypic values. Left, DTF, Pearson’s r, all lines r = 0.77, p < 2.2e-16, df = 108, KO only r = 0.81, p = 5.697e-10, df = 36). Middle, RLN, all lines r = 0.06, p = 0.58, df = 96; KO only r = 0.35, p = 0.037, df = 32. Right, CLN, all lines r = -0.017, p = 0.87, df = 96; KO only r = 0.28, p = 0.11, df = 32). Simple linear model values: multiple r^2^adj[p], log2(DTF ratio KO) ∼ log(*FLC*Wildtype): 0.30 [0.0002], log2(RLN ratio KO) ∼ log(*FLC* wt): 0.18 [0.0068], log2(CLN ratio KO) ∼ log(*FLC*Wildtype): 0.02 [>0.05]). **D**. Correlation between vernalization sensitivity (log2) after eight weeks of vernalization and *FLC* expression levels before vernalization. Black dashed horizontal line indicates Col-0. Mutants: log2(vernalization sensitivity DTF): mean [±sd] 0.23 [±0.21], range -0.20 to 0.82; log2(vernalization sensitivity RLN): mean [±sd] 0.40 [±0.40], range -0.42 - 1.42; wild types: log2(vernalization sensitivity DTF): mean [±sd] 1.41 [±0.49], range 0.17 - 2.41; log2(vernalization sensitivity RLN): mean [±sd] 1.57 [±0.35], range 0.59 to 2.26; KO mutants only: log2(vernalization sensitivity DTF); mean [±sd] = 0.12 [±0.19]), range: -0.20 - 0.5; log2(vernalization sensitivity RLN): mean [±sd] = 0.20 [±0.38]), range: and -0.42 to 0.97. The distribution of vernalization sensitivity, as shown by the histograms, was analyzed separately for KO and KD populations. **E**. Pearson’s correlation coefficients of flowering traits before and after vernalization. *, p ≤ 0.05; **, p ≤ 0.01; ***, p ≤ 0.001; n.s., not significant.

We conclude that mutation of *FLC* accelerates flowering time in all genetic backgrounds tested, although the balance between the vegetative and reproductive portion of the main inflorescence was not always changed in the same manner.

#### Background-dependent action of *FLC*

Compared to their respective wild types, all *flc* mutants flowered earlier, as measured by chronological time (DTF) or developmental time (RLN) (two sided Student’s t test, Benjamini-Hochberg correction, p.adj. < 0.05) (Fig. S1). To estimate how the genetic backgrounds affect the relative contribution of *FLC* to delaying flowering in each genetic background, we asked how well flowering time traits were correlated between *flc* mutants and corresponding wild types. Our first observation was that the relative acceleration in flowering time (the ratio of flowering time between wild type and the corresponding *flc* mutant) was positively correlated with DTF of the wild type, largely irrespective of whether *FLC* was completely or partially inactivated in the mutant. This was more weakly the case for RLN, which suggests that *FLC* affects not only time to bolting, but also the speed with which rosette leaves are produced, i.e., the rosette leaf initiation rate (plastochron). The weakest correlation was seen for CLN, with ratios for KD alleles not being correlated at all (Fig. 1C). Only 83 *flc* lines showed a change in CLN (p.adj. < 0.05). Most, 78 *flc* lines, had fewer CLN than the corresponding wild type, but five mutants representing three accessions (5741, 9102, 9569) showed the opposite, a significant, albeit small increase in CLN, indicating that the effect of the mutation was inverted depending on the genetic background (Fig. 1C and Fig. S2).

Taken together, the effects of *flc* mutations strongly vary across genetic backgrounds and the relative contribution of *FLC* to flowering time traits was least obvious in mutants with KD alleles.

#### *FLC*-independent response to vernalization

*FLC* is the major regulator of the vernalization pathway but much less is known about *FLC*-independent control of vernalization in *A. thaliana* (Michaels and Amasino 1999; Alexandre and Hennig 2008; Deng et al. 2011; Sánchez-Bermejo et al. 2012). For a first glimpse of it, we measured the vernalization sensitivity of 107 *flc* lines and 54 corresponding wild types for DTF, and 91 mutants and 36 wild types for RLN. The *flc* lines were much less sensitive to vernalization than the wild types highlighting the expected strong contribution of *FLC* (Mann-Whitney-U-test [MWU] mutants versus wild types for both DTF and RLN, p < 0.0001). Residual vernalization sensitivities (expressed as log_2_ ratio of flowering time without and with vernalization) of the *flc* KO lines (log_2_[DTF] range -0.20 to 0.54) were on average smaller than of the KD lines (log_2_[DTF] range 0 to 0.82) (Mann-Whitney U rank test, KO versus KD: log_2_[DTF]: p = 1.5e-05; log_2_[RLN]: p = 0.0007). The differences in vernalization sensitivity between the KO and KD population highlights the extent to which vernalization sensitivity can be tuned by modifying *FLC* transcript abundance (Fig. 1D and Table S1), while the residual variation in the KO lines presents an opportunity to dissect the basis of natural variation in the factors controlling *FLC*-independent vernalization.

#### Canalization of flowering traits in *flc* mutants

Across *A. thaliana* accessions, the time of bolting and the time to initiation of the first flower tend to be correlated. This translates into DTF and RLN on the one hand and CLN on the other hand usually change in lock step, but this link can be genetically uncoupled. Differences between accessions with contrasting life history strategies, of which a major determinant is *FLC*, have been reported, but the role of *FLC* in this is largely unclear (Koornneef et al. 1991; Pouteau and Albertini 2009; Salomé et al. 2011; Kinmonth-Schultz et al. 2021). We tested the contribution of *FLC* to linking DTF/RLN and CLN before and after vernalization, which greatly reduces *FLC* activity, by comparing correlations between the three flowering time traits in *flc* mutants and their parental wild types. In the *flc* mutants, DTF, RLN and CLN were highly positively correlated between vernalized and unvernalized treatments (range of Pearson’s *r*: 0.67 to 0.95) (Fig. 1E and Fig. S4). In the unvernalized wild types, trait correlations were much lower (range of Pearson’s *r*: 0.13 to 0.44), but this difference to the *flc* mutants largely disappeared after vernalization (Pearson’s *r* : 0.68 to 0.85). This observation suggests that *FLC* is a major component of trait de-canalization (uncoupling) of flowering traits across natural genetic backgrounds and that vernalization efficiently restores canalization through repression of *FLC*.

### The genetic architecture of *FLC*-independent and extremely early flowering

As *FLC* is a potent floral repressor, its inactivation greatly accelerates flowering (Fig. 1), which raises the question whether there is selection against combinations of *flc* mutations with early-flowering alleles at any of the other genes of the complex network regulating flowering (Blümel et al. 2015; Bouché et al. 2016). QTL mapping of flowering time variation in *A. thaliana* has predominantly involved accessions with winter-annual or intermediate life history strategies, with only few crosses conducted between naturally *FLC* deficient summer-annual accessions (Alonso-Blanco et al. 1998; O’Neill et al. 2008; Simon et al. 2008; Schwartz et al. 2009; Atwell et al. 2010; Li et al. 2010; Salomé et al. 2011; Strange et al. 2011; Grillo et al. 2013; Huang et al. 2013; Dittmar et al. 2014; Sasaki et al. 2015). Our extensive collection of *flc* mutants, which could be considered artificial summer-annual lines, provides an opportunity for deepening our understanding to the basis of *FLC*-independent flowering control. Will we merely replicate previously mapped QTL, or will the complete removal of *FLC* uncover previously hidden or obscured aspects of the genetic architectures governing natural *FLC*-independent flowering time variation?

#### Little transgressive segregation for flowering traits in *flc* mutants

For systematic identification of loci that affect flowering in the absence of *FLC*, we generated 13 segregating F_2_ mapping populations by intercrossing 20 *flc* KO mutants, representing 16 accessions and chosen to provide contrasts in flowering time (overall phenotypic range of 14.1 to 38.0 DTF and 4.8 to 32.6 RLN). All together, they represent the full phenotypic flowering time spectrum of our collection of *flc* mutants (Fig. 1A, Fig. S5, and Table S3). We obtained measurements for DTF, RLN, CLN, and total leaf number (TLN, the sum of RLN and CLN) for 1,829 F_2_ plants (98 to 155 plants per F_2_ population. F_2_ plants flowered between 12 and 42 DTF, 5 to 38 RLN, and 0 to 17 CLN, Fig. 2A and Fig. S6). The phenotypic range of the parents well explained the phenotypic range of the F_2_ populations for DTF and RLN, for most populations, and transgressive segregation was only observed in a few populations (Fig. 2B). This stands in contrast to the rampant transgressive segregation seen in F_2_ populations with segregating functional *FLC* alleles (Salomé et al. 2011). Transgressive segregation was trait-specific, and CLN was uncorrelated with other flowering traits. Flowering trait correlations were very variable in the F_2_ populations (Fig. 2C and Fig. S7), in contrast to the high correlations seen in the parental *flc* mutants (Fig. 1E), indicating that trait canalization due to *FLC* disruption became uncoupled in specific recombinant backgrounds.

**Figure 2:**
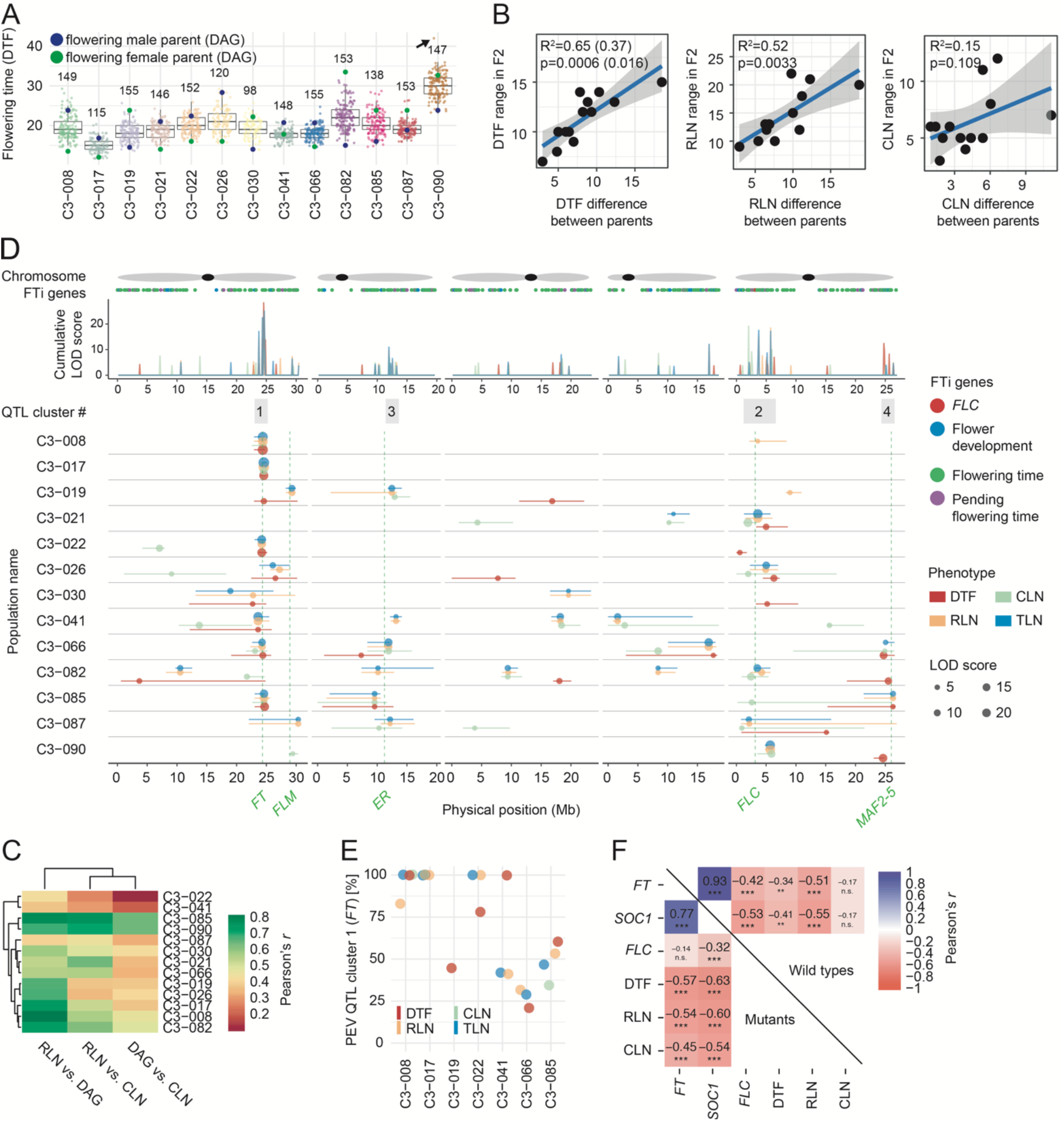
Quantitative genetic analysis of *FLC*-independent flowering. **A**. Distribution of flowering time (DTF) of F2 individuals, including the mean value of the respective mutant parent lines. The number of F2 individuals per population is shown on top. The arrow indicates a single outlier of population C3-090, which was excluded in the correlation shown in C. **B**. Correlation between the phenotypic range of the parents and of the F2 populations (range = F2(max) - F2(min)). Simple linear model, DTF, excluding a single outlier of population C3-090 as indicated by the arrow in Fig 3B: r^2^adj=0.65, p=0.0006; including the outlier: r^2^adj = 0.37, p=0.016; RLN, r^2^adj = 0.52, p = 0.0033; CLN, r^2^adj = 0.15, p = 0.109. **C**. Flowering trait correlations in F^2^ populations. Range of Pearson’s *r*: RLN versus DTF 0.41 to 0.85; RLN versus CLN 0.31 to 0.80; DTF versus CLN 0.13 to 0.68; all p < 0.0001. **D**. Summary of QTL analysis. On top, schematic representation of chromosomes, with black dots representing centromeres. The physical position of genes with a known role in flowering is shown below, colored depending on the flowering category (Blümel et al. 2015; Bouché et al. 2016). LOD scores were summed over a non-overlapping moving window of size 100 kb and shown at the center of the window. The detected QTL clusters 1 to 4 are indicated on top, the widths of the gray boxes indicate the size of each cluster. Cluster 1 is a 2 Mb region (+/- 1 Mb from QTL LOD peak at 24.675 Mb). Cluster 2 is a 7 Mb region (0 to 7 Mbp) on chromosome 5. Clusters 3 and 4 are smaller and contain fewer QTL. All detected QTL in all populations and phenotypic effects are shown below. The QTL intervals (95% Bayes interval) are shown as horizontal lines, and the physical positions of *a priori* flowering candidate genes are indicated as dashed vertical green lines. **E**. Proportional explained additive variation (PEV) of QTL colocalizing with *FT*. **F**. Pearson’s correlation coefficients of expression levels of *FT* and *SOC1* with flowering traits. Mutants on the lower and wild types on the upper triangle. Pearson’s *r*, mutants, -0.57 to -0.45, p < 0.001, 109 d.f.; wild types, DTF: *r* = -0.34, p < 0.01, RLN *r* = -0.51, p < 0.001, 59 d.f. The significance of the correlation is indicated by *, p ≤ 0.05; **, p ≤ 0.01; ***, p ≤ 0.001; n.s., not significant.

#### Four major QTL clusters for *FLC*-independent flowering

Which genes regulate flowering in the absence of *FLC*? To map flowering-related QTL in our *flc* F_2_ populations, we first identified informative genetic markers that distinguished the pairs of parental accessions. To this end, we improved on a method for genome reconstruction and crossover prediction (TIGER, Rowan et al. 2015). This exercise resulted in 627 to 2,648 genetic markers (84 to 694 per chromosome) per accession pair, reflecting the variation in genetic distance between the parental accessions (Table S4).

We identified up to five QTL per population, for a total of 115 additive QTLs, with an average of 2.25 QTL per population * phenotype combination. Only for CLN in population C3-030 were we unable to detect a QTL (Fig. S8A and Table S6).

The total explained additive variation across all QTL ranged from 10% to 66% (Fig. S8B). Most QTL, 99 out of 115, had positive effects, consistent with flowering-promoting alleles contributed by the early parent (Fig. S8C). When summed up per population, all QTL effects were positive and predictive for the differences in flowering time between parental accessions (simple linear model, multiple R^2^_adj_ = 0.68, p = 0.0005) (Fig. S8D and E). Two-dimensional genome scans revealed only three spurious, minor-effect QTL interactions. The lack of epistasis stood in stark contrast to crosses with fully functional *FLC* alleles, indicating that additive flowering-time QTL become prevalent once *FLC* is inactivated (El-Lithy 2005; Simon et al. 2008; Li et al. 2010; Salomé et al. 2011; Strange et al. 2011; Huang et al. 2013; Sasaki et al. 2015) (Table S6). This is in agreement with what has been observed when flowering-time QTL were mapped in populations that were allowed to overwinter outdoors, conditions under which neither *FLC* nor *FRI* make major contributions to flowering time control (Brachi et al. 2010).

To identify shared QTL for each phenotype, we summed up the LOD scores for all F_2_ populations per phenotype in moving 100 kb windows, which allowed us to pinpoint four major QTL clusters. Several narrowly colocalizing QTL defined cluster 1 on chromosome 1, multiple more broadly distributed QTL cluster 2 at the top of chromosome 5, RLN and CLN-specific QTL cluster 3 on chromosome 2, and several QTL, mostly for DTF variation, cluster 4 on chromosome 5 (Fig. 2D).

Cluster 1 on chromosome 1 explained 100% of the proportional explained additive variation (PEV) for all phenotypes of the F_2_ populations C3-008 (parents ID7186-15-R03 [14.3 DTF] and ID9729-01-R05 [29.0 DTF]) and C3-017 (parents ID9908-12-R05 [14.1 DTF] and ID9741-02-R03 [20.9 DTF]), with the only exception being a still high 83% of RLN in C3-008 (Fig. 2D and E). A very strong candidate gene for cluster 1 is the FLC target *FT,* which encodes the long-distance flowering signal florigen (Takagi et al. 2023). We were surprised that the genetic basis of extremely early flowering was that simple - a combination of an engineered *flc* mutation and a natural *FT* allele. Five other populations, which did not have an extremely early flowering mutant as one of the parents, had QTL at cluster 1, explaining PEV from 21% to 100% of at least one phenotype (DTF only in C3-019; DTF, RLN, and TLN in C3-022, C3-041 and C3-066; DTF, RLN, CLN and TLN in C3-085) (Fig. 2D and E).

Most likely, these populations have weaker *FT* alleles, given that we did not find evidence for non-additive suppression of QTL effects. In F_2_ populations C3-019, C3-022, C3-041 and C3-066, we identified additional, non-overlapping QTL, demonstrating that the complexity of the genetic architecture of flowering time traits depends on genetic backgrounds even after inactivation of *FLC*.

#### A hotspot for *FLC*-independent flowering time regulators near *FLC*

QTLs localizing in a wide region around *FLC* have been identified in several studies, and for most, allelic variation at *FLC* itself appears to be the underlying cause. The strong effects of *FLC* obscure the weaker effects of other genes, and it has been difficult to disentangle the contributions of *FLC*-linked loci from those of *FLC* itself in QTL around *FLC* (El-Lithy 2005; Salomé et al. 2011; Strange et al. 2011). Because we analyzed F_2_ populations in a *flc* null background, we could unequivocally test if other loci at the top of chromosome 5 contribute to flowering time differences.

*FLC* is located centrally in the broad cluster 2 composed of 21 QTLs. Together, these QTLs explain variation of all four flowering-time traits in nine F_2_ populations, with a PEV per phenotype from 18% to 100% (average of all populations, DTF 53%, RLN 52%, CLN 46%, TLN 60%) (Fig. 2D and Table S6). Four of the QTLs were located in a narrower region of 1 megabase centered around *FLC*, which explained 18% to 100% PEV in three F_2_ populations (C3-008, RLN 18.4%; C3-021, RLN 100%, TLN 80.5%; C3-082, TLN 33.8%). As *FLC* could be excluded as the cause, we concluded that the top of chromosome 5 is a general hotspot for vernalization-dependent and -independent flowering time regulators (Fig. 2D and Table S6). It will be interesting to determine whether genetic linkage of multiple flowering regulators is advantageous. We note that there are no obvious abnormalities in species-wide linkage disequilibrium in this region, but the known allelic heterogeneity of *FLC* alleles might obscure such patterns (Kim et al. 2007; Salomé et al. 2012; Choi et al. 2013; Li et al. 2014; Baduel et al. 2021). In summary, our findings indicate that *FLC*-independent flowering is governed by QTL clusters that overlap only partly with previously identified QTL. Notably, cluster 1, colocalizing with *FT*, explained significantly more variation as a single QTL than previously observed. Additionally, the expansive cluster 2, around *FLC*, comprises several QTL with strong effects that were likely obscured by *FLC* in earlier studies. Many more QTL with small effects distributed on all chromosomes potentially represent yet unmapped QTL.

#### Uncoupling of flowering traits in F_2_ populations

While leaf number traits were overall highly correlated with DTF, and underlying QTL frequently colocalized, independent QTL contributed to DTF and leaf number traits in three F_2_ populations (C3-019, C3-082, and C3-090) (Fig. 2C and D). Similarly, the genetic basis of leaf number traits RLN and CLN was often shared, but the F_2_ populations C3-022 and C3-026 had also separate QTL for RLN and CLN (Fig. 2D and Table S6), despite comparatively high correlation between RLN and CLN (C3-022, Pearson’s *r* = 0.31 and 0.39, Fig. 2C). F_2_ populations C3-082 and C3-090, which shared ID941-04-R02 as parent, had QTL on top and bottom of chromosome 5, which both contributed to DTF and leaf number but in opposite directions (Fig. 2D and Fig. S8C).

The QTL cluster 4 at the bottom of chromosome 5 (23 - 27 Mb) explained 16% to 100% PEV of RLN and of CLN in one population each, of TLN in two populations, and of DTF in four populations, indicating that this region affects both flowering time and leaf initiation rate, which is related to the ratio of the number of leaves produced before bolting, which is close to TLN, and days to flowering.

The average PEV per phenotype ranged from 17% (CLN) to 51% (DTF). The opposite was found for cluster 3 on chromosome 2: two to four QTL in five populations explained 6% to 100% PEV, especially for CLN (42%), but also RLN (22%) and TLN (26%), but it only contributed to DTF with 9% PEV. QTL specific to CLN were found on chromosomes 1, 3, and 5 (Fig. 2D).

We conclude that genetic coupling of flowering traits is common but uncoupling of specific traits in specific genetic backgrounds could be observed.

### A simple explanation for the extremely early flowering and its unnoticed occurrence in southern Italy

#### Contribution of *FLOWERING LOCUS T* (*FT*) to extremely early flowering

We had concluded that in at least two F_2_ populations, extremely early flowering was triggered by a combination of only two major genetic determinants, the CRISPR-engineered *flc* KO, and a naturally strongly active allele of *FT*. Natural variation at *FT* is widespread and variation in promoter length, and upstream and downstream non-coding regulatory regions are contributing to *FT* expression and adaptation (Schwartz et al. 2009; Adrian et al. 2010; Tiwari et al. 2010; Cao et al. 2014; Liu et al. 2014; Zicola et al. 2019; Takagi et al. 2023). Further, consistent with high levels of *FT* RNA expression being indicative of strong flowering-promoting activity, relative *FT* transcript levels correlated with all flowering time traits in the *flc* mutants and corresponding wild types, except for CLN in the wild types (Fig. 2F and Fig. S2) (Suter et al. 2014; Kinmonth-Schultz et al. 2021).

When compared to the respective wild type, 30 *flc* mutants representing 25 accessions showed increased levels of *FT* (two sided Student’s t test, Benjamini-Hochberg correction, p.adj. < 0.05). *SOC1*, another direct target of FLC and central flowering time integrator, had increased transcript levels in 56 mutants representing 40 accessions (two sided Student’s t test, Benjamini-Hochberg correction, p.adj. < 0.05) (Hepworth et al. 2002) (Fig. S3). Correlations of *SOC1* levels with flowering time traits were very similar to those observed for *FT* (Fig. 2F and Fig. S4). As no QTL colocalized with *SOC1* in our analysis, we reasoned that *SOC1* only mediates the effects of other, polymorphic flowering time regulators (Fig. 2D). It is therefore very likely that the derepression of *FT* by *FLC* in combination with a strong *FT* allele resulted in high *FT* transcript levels that directly induced extremely early flowering.

#### Extremely early-flowering individuals in southern Italy

Since reduction of *FLC* flowering-repressing activity, either through *FRI* or *FLC* mutations, is common in natural populations of *A. thaliana*, and since alleles with high *FT* flowering-promoting activity are common as well, we were wondering why a literature survey suggested that natural accessions that flower as early as our artificial material does in controlled conditions are rare (Lempe et al. 2005; Shindo et al. 2005; Werner et al. 2005; Brachi et al. 2010; 1001 Genomes Consortium 2016; Tabas-Madrid et al. 2018; Fulgione et al. 2022). We confirmed this experimentally by comparison of the earliest flowering accessions (selected from publicly available datasets) to our *flc* mutants (Fig. S11A). We imagined two scenarios to explain this potential paradox: We had created combinations of *FLC* and *FT* alleles that were deleterious in natural conditions, or extremely early flowering individuals do exist in nature but have been mostly missed by collection efforts.

To test the first hypothesis we determined if the *FT* alleles found in extremely early flowering mutants are exclusively present in accessions with a rapid cycling life history strategy. We performed phylogenetic analysis with *FT* sequences of 1,135 accessions and integrated *FLC* expression and flowering data as indicators of the category of life history strategy of different accessions (Atwell et al. 2010; 1001 Genomes Consortium 2016; Kawakatsu et al. 2016). Focusing on the early flowering parents of the five F_2_ populations with a flowering time-related QTL at *FT*, we found that their *FT* sequences fell into different phylogenetic clades. These claded were represented by both summer- and winter-annual accessions (Fig. 3A). Hence, as combinations of *FLC* and *FT* alleles similar to those present in our *flc* mutants are present in natural conditions they are likely not deleterious.

**Figure 3:**
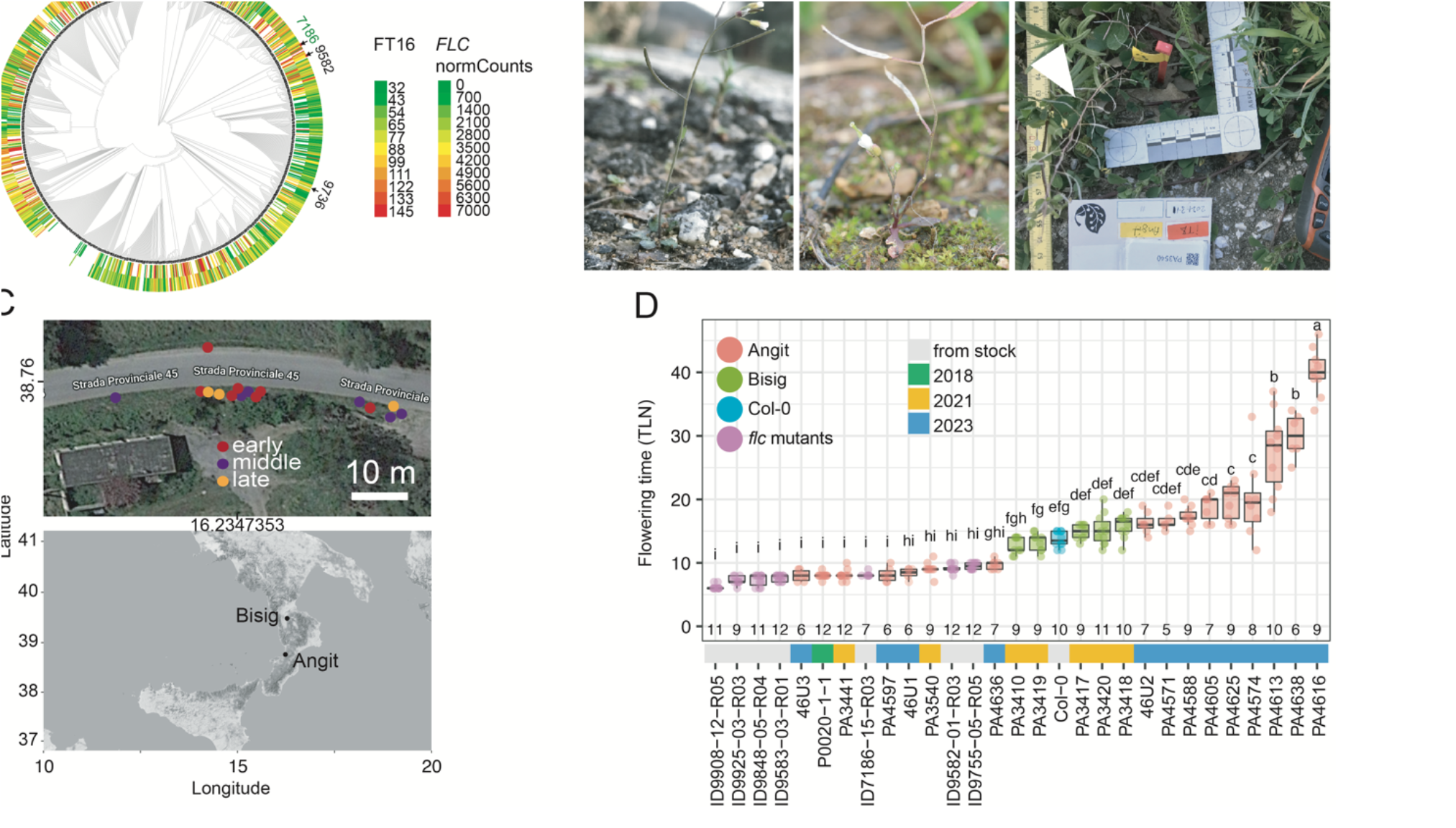
Phylogenetic analysis of *FT* sequences and collection of early flowering samples in Italy. Phylogenetic analysis of *FT* including 382 positions (entire fragment 24,325,373 to 24,335,992 bp of chromosome 1) of 1,135 accessions. The branch lengths of the circular tree were not displayed for easier visualization. **B**. Representative photos of wild plants at the time of sampling at the Angit site. Left: 4636; middle: 4580; right: 3540. **C**. Sampling position at the Angit site (Google Maps) and geographic origin of the Italian sampling sites. Colors indicate the flowering time classified in three groups: Red: early; purple: middle; orange: late. **D**. Flowering time analysis (total leaf number - TLN) at a constant temperature of 22°C and long-day conditions (16 hr light / 8 hr dark) of Italian samples, *flc* mutants, and Col-0. Similar letters indicate no significant difference in total leaf number (ANOVA with post hoc Tukey HSD, p.adj. < 0.05).

To test the second hypothesis, that extremely early flowering individuals do exist in nature but had been mostly missed by previous collection efforts, we speculated that extremely early flowering accessions would most likely have evolved in regions at the hot and dry environmental limit of the species, for example, around the Mediterranean (Putra et al. 2023). We therefore explored locations close to Angitola (Angit) and Bisignano (Bisig) in southern Italy, where we collected a total of 21 early-flowering plants from two populations in 2018, 2021 and 2023 (Fig. 3B, C, and Fig. S11B to K). The progeny of the wild-collected plants from Bisig all flowered moderately early in our greenhouse conditions, but Angit progeny had a range of flowering times, from moderately late to very early (Fig. 3D). Seven of 16 Angit lines flowered as early as the earliest *flc* mutants. These seven lines came from all three collection years, suggesting that a broad range of flowering, including very early flowering is maintained and common in this population (ANOVA with post hoc Tukey HSD, p.adj. < 0.05) (Fig. 3D). Given the very high plasticity of flowering time traits in *A. thaliana*, it was not surprising that the observed flowering time (RLN) in the field did not predict phenotypes obtained in controlled conditions very well (simple lm, adj. R^2^: 0.1547, p > 0.05). Nevertheless, five of 16 (29%) accessions flowered similarly early (Fig. S11L).

Taken together, we found that new mutant phenotypes created in the laboratory may point to portions of the natural phenotypic spectrum of wild plants that are missed because these phenotypes are simply not expected by collectors, in this case, extremely early flowering. We conclude furthermore that extremely early flowering in specific genetic backgrounds can apparently be achieved by changes at only two loci, *FT* and *FLC*.

### Broad pleiotropic roles of *FLC*

#### Contrasting effects of *FLC* mutations on vegetative biomass

Ecophysiologically relevant traits of plants such as growth, biomass accumulation, and leaf structure are often correlated, forming a trade-off known as the leaf economics spectrum (LES) (Wright et al. 2004; Sartori et al. 2019). Contrasting solutions to this trade-off are associated with specific life history strategies, such as differences in flowering time (Wright et al. 2004; Sartori et al. 2019). The so-called slow-fast-continuum, as prevalent in many plant species, refers to the co-occurrence of either earlier flowering and faster growth, or later flowering and slower growth, and it is tightly linked to the leaf economics spectrum.

In addition to flowering, *FLC* is a regulator of another post-embryonic developmental transition, the switch from the juvenile to the adult vegetative phase (Wu 2006; Schwarz et al. 2008; Mentzer et al. 2010; Willmann and Poethig 2011). Whilst most differences in leaf morphology caused by changes in *FLC* activity appear to be indirect through the *FLC* effects on flowering, the impact of *FLC* on leaf shape appears to be flowering-independent (Willmann and Poethig 2011). To test whether *FLC* affects the growth rate or consistent differences in plant size, and whether the extent of an effect is dependent on the genetic background, we measured the projected rosette area, as retrieved from top-view images (Sartori et al. 2019), over time. It serves as a proxy for biomass, which in turn is a proxy for fitness (Younginger et al. 2017; Vasseur et al. 2018a). From the previously described experiment (Fig. 1A), we selected for measurements the time period of 11 to 18 days after germination, which corresponds for most lines to the phase of early vegetative biomass accumulation. As described earlier, five *flc* mutant lines representing three accessions already bolted 16 days after germination. However, analysis of growth trajectories under comparable growth conditions has revealed that growth only slows down one week after bolting (Hanemian et al. 2020), therefore no confounding effect of even extremely early flowering on growth was expected in the time frame selected for growth measurements.

An analysis of the relative growth rate (RGR) between day 11 and 18 revealed no significant differences between the pairs of mutants and wild types (Mann-Whitney U rank test, Benjamini-Hochberg correction, p.adj. > 0.05) (Fig. S12A and B). Then, we focused on consistent size differences on each day of measurement. Overall, the projected rosette area of the *flc* lines and the wild types were similar, regardless of the day of measurement (Mann-Whitney-U-test, Bonferroni correction, p.adj. > 0.05) (Fig. S12C) (Vanhaeren et al. 2015). Seventy-seven mutants did not show a size difference at any of the timepoints (Mann-Whitney-U-test rank test, Benjamini-Hochberg correction, p.adj. > 0.05). We focused on the remaining 33 *flc* mutants representing 27 accessions that were different from the corresponding wild type on at least one day (Fig. 4A). We clustered these mutants using the ratio of mutant versus wild-type projected rosette area, and found three major clusters. The 16 *flc* mutants representing 13 accessions in cluster 1 were consistently larger than the respective wild type. Three *flc* mutants representing three accessions in cluster 2 were slightly larger on only one or two days. In other words, cluster 1 and 2 consisted of *flc* mutants that followed the slow-fast-continuum. The most interesting cluster, cluster 3, included 14 *flc* mutants representing 11 accessions that were consistently smaller than the corresponding wild type (Mann-Whitney U rank test, Benjamini-Hochberg correction, p.adj. < 0.05) (Fig. 4A and Fig. S12D). That both cluster 1 and 3 included multiple mutants from the same accession, without any overlap in accessions between these two major clusters, supports that the observed patterns are due to *FLC* mutations.

**Figure 4:**
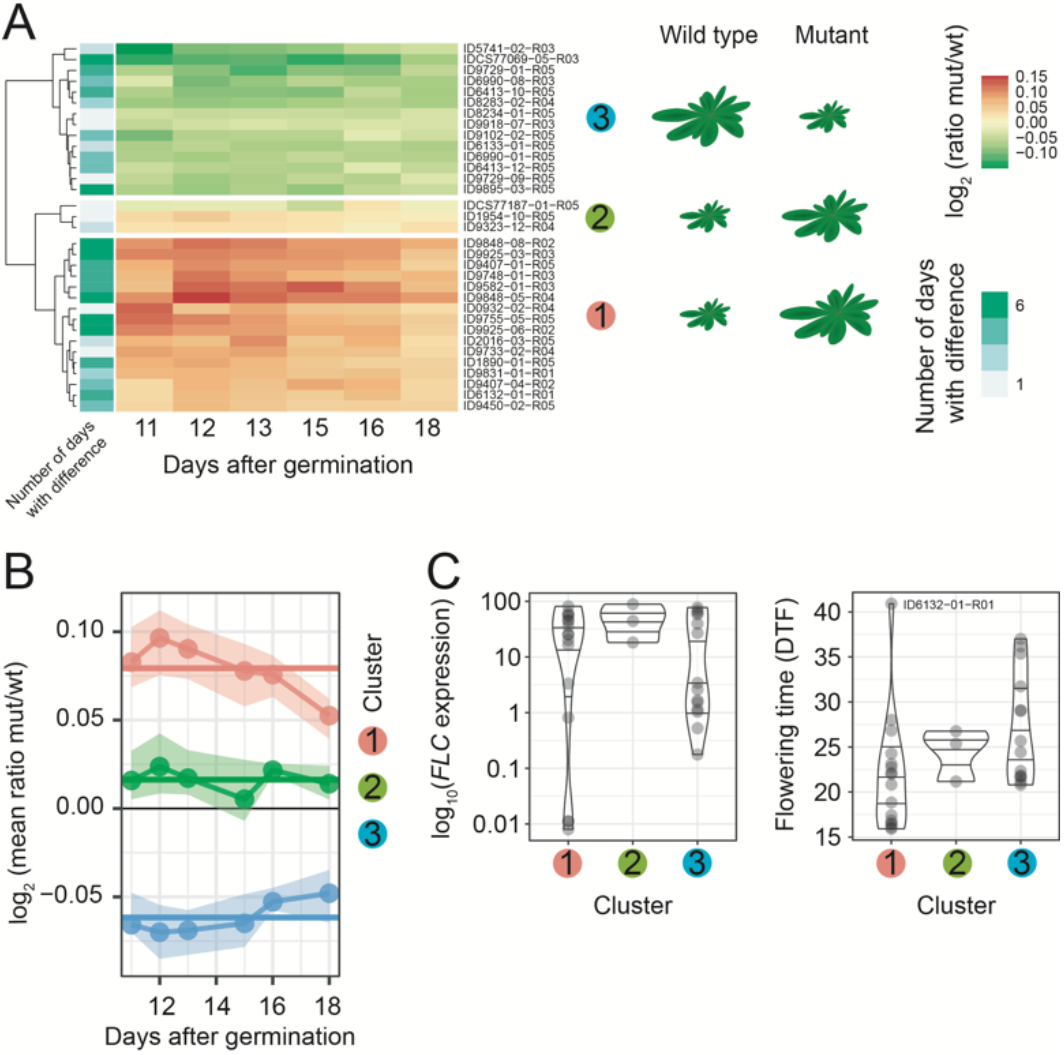
Analysis of wild-type and mutant growth trajectories. **A.** Clustering was performed using the projected rosette area ratio (log2) of mutants versus wild types, with a total of 110 mutant lines included. The projected rosette area of the wildtype 8376 could not be determined, so the lines ID8376-04-R05 and ID8376-10-R04 were excluded. The number of days with significantly different projected rosette area measurements (Mann-Whitney-U-test, Benjamini-Hochberg correction, p.adj. > 0.05) is indicated. A schematic representation shows the cluster size differences. **B**. Cluster means of the ratio on each day (dots) and over all days (horizontal lines, log2). The values between 25^th^ and 75^th^ percentiles are shown as ribbons. **C**. *FLC* levels (log) and flowering time of mutants in cluster 1 to 3.

The accessions in cluster 1 and 3, which were either consistently larger or smaller than the corresponding wild types, were genetically diverse and belonged to six and seven different genetic groups as determined by (1001 Genomes Consortium 2016), with shared membership in six groups. The average projected rosette area ratio per cluster was constant over the selected time period for all clusters (Fig. 4B). No differences in flowering time (DTF) and *FLC* transcript levels were observed between the mutants of the three clusters (Kruskal-Wallis H-test, p > 0.05), indicating that differences in *FLC* levels and flowering time alone do not explain the relative growth differences (Fig. 4C). However, all the earliest flowering mutants (DTF < 20) belonged to the large cluster 1, showing that at the early flowering end of the phenotypic spectrum, the lines strictly followed the slow-fast-continuum, whilst late-flowering lines were part of all clusters.

#### Diverse and background-specific effects on global gene regulation

In addition to the reported roles of *FLC* in controlling major life history transitions, like germination, the juvenile-adult transition, and the vegetative-reproductive transition, *FLC* likely has additional regulatory roles as suggested by its binding to hundreds of target genes with functions in many different developmental pathways (Deng et al. 2011; Mateos et al. 2015).

To reveal potential background-specific regulatory roles of *FLC*, we performed a RNA-seq analysis of seven early flowering *flc* mutants (DTF < 30), with low *FLC* transcript levels (*FLC* qPCR level range 0.1 to 11.7 a.u., all KOs except ID9402-01-R05 = 11.7 a.u.) (Fig. 5A and Table S1). We extracted the differentially expressed genes (DEGs) of each *flc* mutant/wild type contrast (FDR < 0.1 and |log_2_(FoldChange)| > 1). Besides *FLC*, which was downregulated in all contrasts by -8.3 to -4.0 (logFC), we identified a total of 31 unique differentially expressed genes (DEGs). The number of DEGs per contrast excluding *FLC* ranged from 1 to 17. Excluding *FLC*, we found DEGs to be upregulated in 21 cases (mean [±se] logFC 3.0 [±0.38]) and downregulated in 18 cases (mean [±se] logFC -3.3 [±0.43]). Our previous analysis had shown that *SOC1* was upregulated in most *flc* mutants and we detected *SOC1* as differentially expressed in all contrasts (range logFC 2.3 to 5.3), irrespective of the *flc* mutant or wild type flowering time difference (Fig. 5A and Fig. S1C). Besides *SOC1*, only one other flowering-time gene (out of a list of 428 genes, Table S5), *SVP*, was upregulated in two contrasts (logFC 1.8 and 2.4) (Blümel et al. 2015; Bouché et al. 2016). In three contrasts, only *FLC* and *SOC1* were differentially expressed. Of the 28 non-flowering related DEGs, 27 were restricted to one contrast, and one DEG, which is not a known target of *FLC*, was shared between two contrasts (AT5G22580, logFC 1.9 and 2.5). *PAP16*, a known target of FLC, was strongly upregulated in the *flc* mutant ID9729-01-R05 (logFC [FDR]: 7.0 [0.083]) (Deng et al. 2011; Mateos et al. 2015, 2017) (Fig. 5A). Due to the timing of the sampling during the generative phase transition we were not able to distinguish between *FLC* direct regulatory effects and those that are indirect consequences of a change in flowering time in *flc* mutants. Nevertheless, the magnitudes of flowering-time differences were not predictive of DEG sets, indicating that gene regulatory effects of *FLC* mutations and the genetic networks, in which *FLC* participates in, vary between different genetic backgrounds.

**Figure 5:**
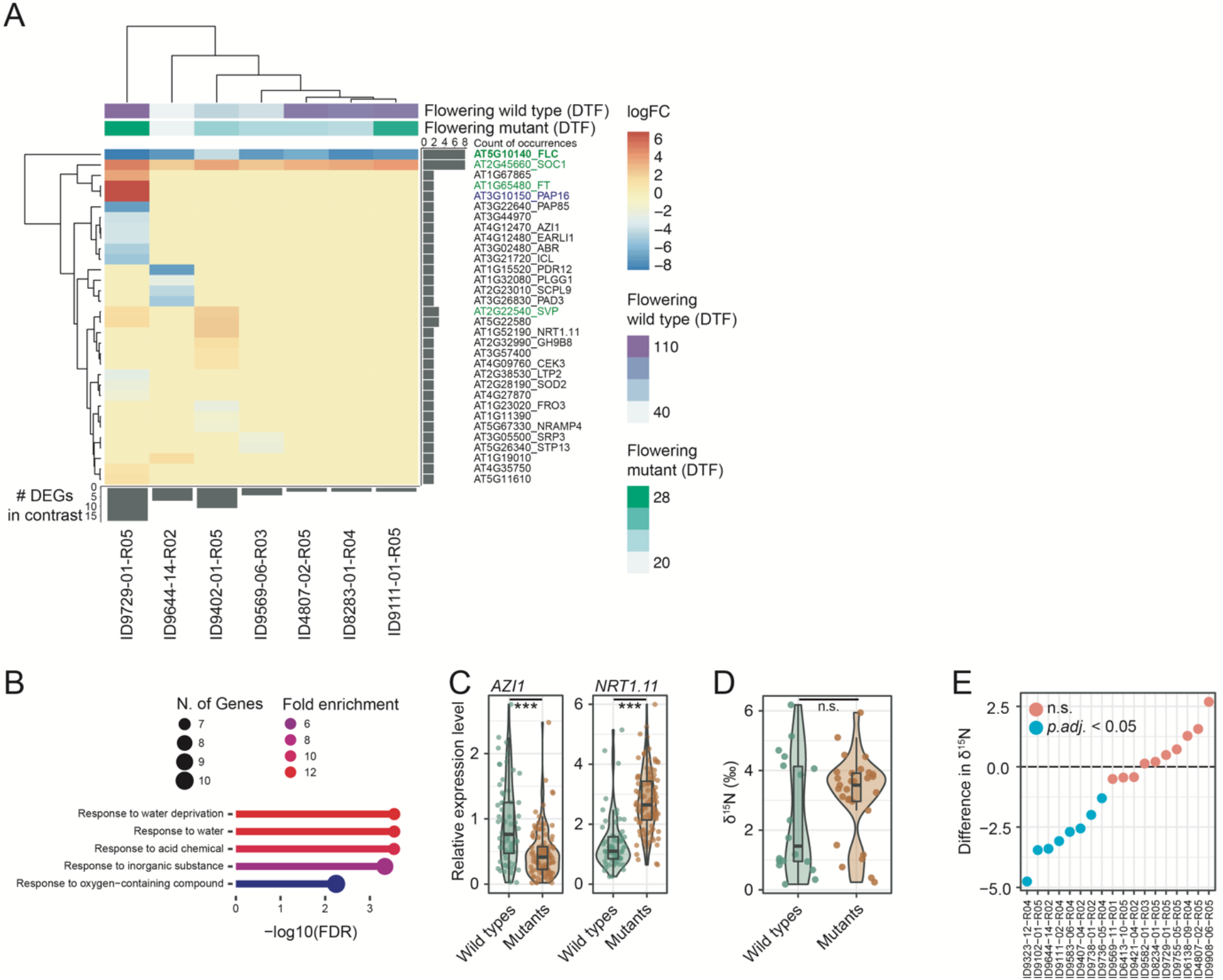
Detailed analysis of *FLC*’s pleiotropic roles. **A**. Heatmap of the logFC values of differentially expressed genes (DEGs) present in at least one contrast. Flowering time data (DTF) from the experiment shown in Fig. 1A. **B**. GO enrichment analysis. The top 5 hits (-log(FDR)) are shown with an FDR cutoff of 0.01. **C**. Nitrogen isotope composition (δ^15^N [‰]) in mutants and wild types (means of three biological replicates). Mean [±se] mutants: 3.17‰ [±0.26]; wild types: 2.43‰ [±0.46]; Mann-Whitney U rank test, p > 0.05. **F.** Relative expression levels [a.u.] of *AZI1* and *NRT1.11* in the wild types and the mutants. *AZI1*, mean wild types: 0.92 a.u., mean mutants: 0.47 a.u., *NRT1.11*, mean wild types: 0.47 a.u., mean mutants: 1.37 a.u. **D.** Nitrogen isotope composition (δ^15^N [‰]) in mutants and wild types (means of three biological replicates). Wild types, N=18; mutants, N=29. Mean [±se] mutants: 3.16‰ [±0.26]; mean wild types: 2.43‰ [±0.46]; Mann-Whitney U rank test, p = 0.17. **E**. Difference in δ^15^N (δ^15^Ndiff = δ^15^NWildtype - δ^15^NMutant) between wild type and the respective mutant. The blue dot indicates a significant difference, two-sided Student’s t-test, Benjamini-Hochberg correction, p.adj. < 0.05.

To identify a potentially common developmental role of all 31 unique DEGs (excluding *FLC*), we asked whether any GO terms were enriched among these DEGs. One of the enriched categories was for water stress related groups (-log(FDR) > 2) (Fig. 5B). This caught our attention, as flowering time shapes drought escape strategies of *A. thaliana* accessions, and the *FRI*/*FLC* module has a pleiotropic role in drought avoidance strategy by regulating water use efficiency (McKay et al. 2003, 2008; Lovell et al. 2013; Easlon et al. 2014; Kenney et al. 2014). Only recently have we demonstrated that the commonly constrained traits water use efficiency and flowering time are uncoupled in specific *flc* mutants (Ruffley et al. 2024).

#### Broad physiological, background-dependent pleiotropic roles of *FLC*

Our transcriptomic analyses of *flc* mutant tissues also pointed to broad pleiotropic roles of *FLC*. In the *flc* mutant of wild type 9729, *AZI1* and *EARLI1* were downregulated (logFC [FDR]: -3.46 [0.039] and - 3.43 [0.083]), two genes with roles in systemic defense priming and control of root-growth under Zn-limiting conditions (Cecchini et al. 2015; Bouain et al. 2018). The *flc* mutants generally had lower levels of *AZI1* than the wild types (Mann-Whitney-U-test, p = 9.4e-08) (Fig. 5C). While the difference in *AZI1* was not robust to correction for multiple comparison (two sided Student’s t test, Benjamini-Hochberg correction, p.adj. > 0.05), it is worth noting that, without multiple testing correction, *AZI1* was only downregulated in twelve *flc* mutants (representing seven wild types) when compared to their respective wild types (two sided Student’s t test, p < 0.05) (Fig. S12A). The *flc* mutant of wild type 9402 showed upregulation (logFC [FDR]: 2.5 [0.025]) of the nitrate transporter gene *NRT1.11*, which is required to transfer root-derived nitrate into phloem in the major veins of mature leaves (Hsu and Tsay 2013; Guan 2017) (Fig. 5A). *flc* mutants generally had higher levels of *NRT1.11* than the wild types (Mann-Whitney-U-test, p = 2.2e-16) (Fig. 5F). Before multiple testing correction, 33 mutants representing 28 wild types showed differential expression of *NRT1.11*, which was upregulated in all but one case (two sided Student’s t test, Benjamini-Hochberg correction, p.adj. < 0.05) (Fig. S12B), of which one *flc* mutant (ID9743-03-R05) tested significant after correction for multiple testing.

Nitrate is known to regulate flowering partially via *FLC* (Lin and Tsay 2017; Teng et al. 2019). Under non-limiting nitrate conditions, the nitrogen isotope ratio (δ^15^N [‰]) can serve as a proxy for *in planta* nitrogen use efficiency (Evans 2001; Kalcsits and Guy 2013; Carlisle et al. 2014; Craine et al. 2015; Guan 2017). We measured δ^15^N in 20 wild-type accessions (including Col-0) and 29 *flc* mutants (with a maximum relative *FLC* expression level of 18.3 a.u.), which included 19 *flc* mutant / wild type contrasts. The wild types had a bimodal δ^15^N distribution, while such a bimodal distribution was not obvious in the *flc* mutants. Overall, the δ^15^N levels in *flc* mutants and wild types were similar (mean [±se] mutants: 3.17‰ [±0.26], wildtypes: 2.43‰ [±0.46], Mann-Whitney U rank test, p > 0.05). However, pairwise comparisons of 18 *flc* mutant/wild type contrasts revealed eight mutants from six genetic groups with higher δ^15^N (two-sided Student’s t test, Benjamini-Hochberg correction, p.adj. < 0.05) (Fig. 5D, E, and Fig. S13A). δ^15^N correlated neither with flowering time nor *FLC* expression nor δ^13^C levels (from Ruffley et al. 2024) (Pearson’s correlation coefficient, DTF: *r* = 0.3; *FLC* expression level: *r* = -0.12; δ^13^C: *r* = 0.089, p > 0.05). The ratio of carbon to nitrogen (C/N) in plant tissue, which is directly linked to photosynthetic activity, was unchanged in the *flc* mutants relative to the corresponding wild types (two sided Student’s t test, Benjamin-Hochberg correction, p.adj. > 0.05) (Fig. S13B) (Otori et al. 2017; Thieme et al. 2022). Together, this suggests that *FLC* has a background-specific pleiotropic effect on δ^15^N, but that this effect is independent from the other investigated traits.

## Discussion

The continuous advancement of CRISPR/Cas9 gene editing technology and its adaptation for use in various plant species has transformed genetic studies. These methods have liberated researchers not only from the constraints of static mutant collections, but also from the use of limited, standardized genetic backgrounds. While there are several examples of CRISPR/Cas9 knockouts of orthologous genes in different species, especially in crops, investigations of similar genetic perturbations in different genetic backgrounds remain relatively rare (Cho et al. 2017b; Kumlehn et al. 2018; Wu et al. 2018; Ursache et al. 2021; Bieluszewski et al. 2022; Srikant et al. 2022; Ahmar et al. 2023). In a previous study, we described a new collection of *flc* mutants in 62 accessions, showing that *FLC* has background-dependent pleiotropic effects on water use efficiency and that uncoupling of the commonly constrained traits flowering time and water use efficiency can be observed in specific genetic backgrounds (Ruffley et al. 2024).

In the present study, we further leveraged this *flc* mutant collection and uncovered a previously overlooked part of the flowering time spectrum in natural accessions as well as coupling between flowering time traits (Fig. 1 and 2). The genetic basis for extremely early flowering was surprisingly simple, with phenotypic counterparts identifiable in natural populations (Fig. 3). Beyond the life-history trait flowering time, we also discovered background-specific roles of *FLC* in early vegetative growth and broad roles in developmental and physiological processes (Fig. 4 and 5).

### *FLC*-independent vernalization response

We found considerable *FLC*-independent variation in the sensitivity to respond to prolonged cold periods (Fig. 1D). Several genes other than *FLC* have been identified to act on vernalization (Alexandre and Hennig 2008; Sánchez-Bermejo et al. 2012), including the *MAF* family of *FLC* homologs, which are, like *FLC,* vernalization-responsive flowering regulators (Sung et al. 2006). We found QTLs explaining flowering time that colocalize with *MAF* genomic regions on chromosome 1 and 5, and allelic variation in *MAF* genes likely contributes to differences in *FLC*-independent vernalization sensitivity (Michaels and Amasino 2001; Alexandre and Hennig 2008). It remains to be tested how *FLC*-independent variation in vernalization sensitivity is expressed under shorter, less saturating vernalization treatments and under shorter, less flowering-promoting photoperiods. Through biparental crossings of *flc* KO mutants with contrasting vernalization sensitivity, it will now be possible to pinpoint the genetic architecture of the observed range in *FLC*-independent vernalization sensitivity.

### Extremely early flowering, its genetic basis, and possible ecological niches

We used cost efficient quantitative genetics at scale to determine the *FLC*-independent genetic architecture of flowering time traits by crossing *flc* KO lines of genetically diverse backgrounds and with contrasting phenotypes. In agreement with the large body of literature on the genetic architectures of natural variation in *A. thaliana* flowering, we found few loci explaining much of the variation, with predominant contribution from additive genetic effects (Fig. 2) (Alonso-Blanco et al. 1998; O’Neill et al. 2008; Simon et al. 2008; Schwartz et al. 2009; Atwell et al. 2010; Li et al. 2010; Salomé et al. 2011; Strange et al. 2011; Grillo et al. 2013; Huang et al. 2013; Dittmar et al. 2014; Sasaki et al. 2015). However, since transgressive segregation was observed for specific traits, the relatively small size of the mapping populations and multiple testing penalties in two-dimensional genome scans might have obscured the detection of interacting QTLs.

We identified a QTL of major effect that colocalizes with the central flowering regulator *FT*, explaining most of the extremely early flowering of some of the *flc* mutants and detectable in more than half of the tested F_2_ populations. As far as the proportional phenotypic contributions are comparable across the variety of published mapping and association experiments and environmental conditions used therein (Alonso-Blanco et al. 1998; O’Neill et al. 2008; Simon et al. 2008; Schwartz et al. 2009; Atwell et al. 2010; Li et al. 2010; Salomé et al. 2011; Strange et al. 2011; Grillo et al. 2013; Huang et al. 2013; Dittmar et al. 2014; Sasaki et al. 2015), our observations of *FT* being the major contributor to flowering time variation are surprising, reflecting the special nature of our material. Previous studies almost never identified a monogenic architecture of flowering with *FT* explaining nearly all of the phenotypic variation, as it is common for *FLC* (Alonso-Blanco et al. 1998; O’Neill et al. 2008; Simon et al. 2008; Schwartz et al. 2009; Atwell et al. 2010; Li et al. 2010; Salomé et al. 2011; Strange et al. 2011; Grillo et al. 2013; Huang et al. 2013; Dittmar et al. 2014; Sasaki et al. 2015). A notable exception was a cross between two naturally early flowering accessions, which led to the report of a flowering time QTL at *FT*, after more than a decade of work on the genetic basis of natural variation in flowering time in *A. thaliana* (Alonso-Blanco et al. 1998; Schwartz et al. 2009).

The *FT* and *SOC1* transcript levels in our *flc* mutants correlated well with flowering time in our experiments. Complex natural variation in promoter length, and upstream and downstream non-coding regulatory regions are contributing to *FT* expression and adaptation, and we found contrasting polymorphisms in mapping populations with a QTL at *FT* (Schwartz et al. 2009; Adrian et al. 2010; Cao et al. 2014; Liu et al. 2014; Zicola et al. 2019). Through phylogenetic comparison of *FT* sequences and life-history classification that considered *FLC* transcript levels and flowering time data of 1,135 accessions we arrived at the hypothesis that the early flowering of some *flc* mutants should reflect a yet undescribed extreme end of the natural phenotypic spectrum of flowering with potential adaptive relevance. Given the relatively small number of accessions and growth conditions covered by our study, we strongly suspect that the lower limit of flowering times in natural accessions has not yet been found.

Earlier flowering is generally under selection throughout the range of *A. thaliana*, and even more so in Italy, where some regions are at the edge of the apparent environmental limit of the species (Hoffmann 2002; Munguía-Rosas et al. 2011; Ågren et al. 2016; Exposito-Alonso et al. 2017; Putra et al. 2023). Sampling bias could be one of the reasons for the underrepresentation of extremely early flowering accessions in *A. thaliana* seed stocks. This could be driven by the fact that such individuals are ephemeral, and can be collected only during a very short time window.

From a reverse ecological perspective and with a focus on this specific, fast cycling life-history strategy, we undertook a targeted collection effort, through which we found extremely early flowering accessions in southern Italy (Fig. 3). It is difficult to predict flowering time in natural conditions from measurements in controlled laboratory conditions because of strong genotype-by-environment interactions (GxE) and because of the highly dynamic changes of relevant parameters such as temperature and light, which are next to impossible to simulate in the lab. It was therefore not surprising that only a fraction of Italian lines sampled here appeared to have similar phenotypes in the wild and the lab. Further experiments in controlled and common garden experiments will determine the phenotypic plasticity and the degree of GxE. Integration of present climate data should hopefully allow predictions of specific ecological niches.

A possible ecological niche for extreme early flowering could be an environment that supports only a very short life span, which is set by the appearance of destructive conditions such as terminal droughts. Early flowering as a common drought escape strategy would then manifest in an extreme way as “super escapees”, that could be advantageous in regions like Southern Scandinavia where the growth season was predicted to be too short to allow an escape of “normal” summer-annual, early flowering accessions (Exposito-Alonso et al. 2017; Dittberner et al. 2018). Several generations per year, each occupying a short time window, are imaginable.

Given the ample genetic opportunities *A. thaliana* has to accelerate flowering and to escape drought, taken together with potential sampling bias, it seems likely that we have engineered a phenocopy of accessions that are part of the naturally occurring phenotypic spectrum. Is the predicted trait response to future climates reality already? Dedicated and geographically diverse re-sampling of natural populations will show this.

### Unmasking genomic hotspots of *FLC*-independent flowering regulation

Apart from a major-effect QTL that co-localizes with *FT*, we have identified numerous QTL explaining *FLC*-independent flowering-related traits. It is difficult to know whether truly new associations were identified, as confidence intervals of QTL can span large chromosomal regions and the QTL identified here often overlap with published QTL intervals. Strong candidate genes for several of our QTL are *FLM*, *ER*, and the *MAF* gene cluster (Salomé et al. 2011). *FRI*, whose function on flowering time regulation depends on *FLC*, did not contribute to flowering time variation in our material, as expected. Remarkably, we identified the region around *FLC* as another hotspot for flowering time variation (Fig. 2E). In contrast to most other studies, our unique design allows us to exclude any contribution from *FLC* itself, as only mutants with complete *FLC* disruptions were crossed to obtain mapping populations. In previous analyses, segregating large-effect epistatic *FLC* alleles and genetic linkage complicated the separation of minor-effect loci in this region (El-Lithy 2005; Salomé et al. 2011; Strange et al. 2011). Our results present yet another example of how mutations in different genetic backgrounds can reveal cryptic fractions of genetic architectures of adaptive traits.

### Trait canalization and a possible genetic basis

Strong canalization of all flowering time traits was observed in the *flc* mutants. Relationships between the traits became stronger in the mutants, with a clear gradient in de-canalization in the wild types that correlated with flowering times and *FLC* expression levels. In agreement, vernalization, which greatly reduces *FLC* activity, led to trait canalization comparable to the effects of the *FLC* mutations. We determined the number of cauline leaves (CLN) as a proxy for the time between initiation of flowers and bolting, which dictates inflorescence architecture critical for optimal seed set and fitness (Ratcliffe et al. 1999; Pouteau and Albertini 2009; Yamaguchi et al. 2014). *FT*, the major contributor to flowering time variation in the absence of *FLC*, as determined in our crosses, has not only a role as central regulator of induction of flowering but also controls fruit number by timing the inflorescence meristem arrest (González-Suárez et al. 2023). Indeed, *FT* transcript levels correlated with flowering time traits and the time until the first flower (CLN) in the mutants. In the wild types, this was only the case with the flowering time traits, but not with CLN, variation which is largely explained by QTL at *FT.* We identified additional QTL for CLN and other flowering time traits across all five chromosomes as well as a CLN-specific QTL on top of chromosome 5 close to *FLC*, which has not been described in previous mapping experiments or was previously masked by *FLC* (Kearsey et al. 2003; Atwell et al. 2010; Huang et al. 2013; Gupta et al. 2020). Hence, CLN variation of the *flc* mutants seems to be mainly controlled by allelic variation of *FT*, with additional contributions from *FT*-dependent and - independent regulators. Surprisingly, the effect of *FLC* mutations on CLN in the different genetic backgrounds was not exclusively negative, as observed for the flowering time traits (Fig. 1C). The accessions in which this was the case both belong to the genetic group of *Western Europe*, and it will be interesting to investigate if a common genetic basis for the inversion of a trait relationship can be identified (1001 Genomes Consortium 2016).

### Exceptions from the general flowering-growth trade-off

Early or late flowering is often part of trait syndromes and co-variation with biomass accumulation is common (Vasseur et al. 2018b; Takou et al. 2019). In our study, many *flc* mutants were different from the corresponding wild type during the early phase of vegetative growth (Fig. 4). We are aware that these size differences could simply be generated by the experimental procedure as differences in germination, another major life history event that is often associated with flowering time in multi-trait syndromes, are leading to different developmental ages and plant sizes at the time of measurement. The parental plants had been propagated under similar growth conditions, but the wild types had been additionally vernalized for eight weeks, which may have led to better germination of the wild-type progeny (Auge et al. 2017; Blair et al. 2017). Further, active *FRI* in a *FLC* functional background or high *FLC* activity, present in some wild types, enhances germination proportion (Chiang et al. 2009; Auge et al. 2017). Taken together, if differences in germination timing would be the driver for rosette size differences in our analysis, the wild types should be overall larger, which was not the case (Fig. 4). Also, effects of germination were reduced by thinning bulks of plants to a single plant at the beginning of the experiment to obtain an experiment-wide matched developmental age of plants.

Most *flc* mutants among the ones that differed in size from the corresponding wild type were larger. Hence, a modification of the life history strategy through mutation of *FLC* mostly led to a change in growth along the slow-fast-continuum. With that they have favored the acquisition over the retention of resources (Wright et al. 2004; Vasseur et al. 2018b; Sartori et al. 2019). However, this could not be generalized, as mutants with inverse relationship of flowering and growth, as was expected from the slow-fast-continuum, were found (represented by cluster 3). Trait syndromes mediate adaptation to local conditions but the co-variation between traits in natural accessions is not fixed, and correlations in natural accessions can change, e.g., along a latitudinal gradient (Debieu et al. 2013; Vasseur et al. 2018b). As life-history traits determine fitness, one explanation for the deviation from the common slow-fast-continuum (cluster 3 mutants) could be the experience of harsh environmental conditions. As a high growth rate decreases drought tolerance due to the extensive gas exchange and water loss, early flowering together with a slow growth could be advantageous in conditions with severe conditions of drought (Metcalf and Mitchell-Olds 2009; Debieu et al. 2013). Further, staying small in size could be beneficial under conditions of exceptionally high pathogenic load, as expression of immunity genes increases with the length of the vegetative lifespan, leaving early-flowering plants more unprotected (Glander et al. 2018).

While the number of accessions included in our study is still too small to answer these questions definitively, we have demonstrated how the modification of one trait that is part of a complex trait syndrome, through engineering of a central regulatory gene, can unmask deviations from general trait correlations. These “outlier situations” are particularly interesting candidates for functional follow-up studies, as they might provide explanations on how genetic correlations are restructured to adapt to niche environments.

### Broad pleiotropic roles of *FLC*?

While we are aware that our transcriptomic analyses have limitations, as they included only seven mutant/wild type contrasts and a single environmental condition, the data already provide many indications for intraspecific variation of gene regulatory effects and pleiotropic roles of *FLC*. Since the plants for the transcriptome studies had been grown under inductive long-day conditions, we can not clearly distinguish between direct transcriptional effects of *FLC* and indirect effects due to altered flowering time. Nevertheless, neither the absolute flowering time of the mutants nor of the parental wild types nor the relative effect of *FLC* on flowering time traits explain the main differences in differential expression patterns, suggesting that most effects were related to the specific genetic backgrounds. Orthologous analysis of *FLC* binding sites in the genomes of *A. thaliana* and its relative *Arabis alpina* have revealed that fewer than one in five are conserved. Many of the nonconserved target genes are involved in stress responses (Mateos et al. 2017). Likewise, we find that nearly all shared DEGs in our mutant/wild type contrasts are flowering time genes, while stress-related genes are overrepresented among the DEGs that are specific to subsets of accessions or even individual accessions. Moving forward, it would be interesting to also perform transcriptomic analysis in non-inductive short days to detect flowering-independent effects, and to perform *FLC* ChIP-seq experiments in a subset of genomic backgrounds to determine the intraspecific variation of the *FLC* targetome. Even though multiple testing penalties removed statistical support for most of the initially detected effects of *FLC* mutations on expression levels of other tested candidate genes, the results pointed to *flc* mutations affecting genes in diverse developmental and physiological pathways as a function of the genetic background. With our work, we have provided a first glimpse into the broad impact that the central flowering-time regulator *FLC* might have on different physiological levels.

Finally, we would like to emphasize that background-dependent pleiotropic effects are a major limitation in breeding. Simultaneous mutation of the same gene in many different backgrounds and comprehensive analysis provides an excellent avenue for understanding the true extent of genetic networks in which a focal gene participates and such approaches could generally help to engineer quantitative trait variation that goes beyond what is observed in the original population.

## Material and Methods

### Genetic resources

#### *flc* mutants

Described in Ruffley et al. 2024.

#### Generation of F_2_ mapping populations

KO mutant lines covering most of the flowering time range in the mutant population were crossed to generate F_1_ progeny. Per cross, around five F_1_ plants were selfed to obtain biparental F_2_ mapping populations, from which one was randomly selected for the mapping experiment. All plants were propagated at 22°C under long days (LD, 16 h light / 8 h dark). The identity of the parents of the mutant lines and of the F_1_ individuals was verified with shallow Illumina short-read whole-genome sequencing (WGS) and SNPmatch (version 5.0.1) (Pisupati et al. 2017).

#### Plant collections in Southern Italy

Plant populations were identified in 2018 as part of the Pathodopsis collection (Karasov et al. 2022), and based on their proximity to the coordinates listed as original collection sites of accessions from the 1001 Genomes Project (1001genomes.org): 38.76°N, 16.24°E (Angit) and 39.48°N, 16.28°E (Bisig). In 2018, a single plant (P0020-1) that was already mature in the Angit population at the time of visit was collected in a seed bag. In 2021 and 2023, plant collections followed a linear transect through the entire extent of both populations, and roughly every 20th plant encountered was selected for seed propagation. The precise coordinates of the plants harvested for this study are listed in Table S7. Prior to harvesting, plants were photographed with an iPad pro, usually including a photomacrographic scale ABFO No.2 (cop-shop.de) as size reference. Collection spots were either directly georeferenced with a GPS (Garmin GPSmag 64S) or indirectly by distance to a georeferenced point along the transect. Mature plants were then directly transferred into a seed bag, less mature plants were transferred, with their roots, into pots with potting soil (CL P, einheitserde.de). After transport back to Germany by car, plants in pots were grown on a windowsill until seeds could be harvested. From every seed stock obtained in 2018 and 2021, a single individual was propagated in a growth chamber in the laboratory at 23°C under LDs to obtain a fresh seed stock. In 2023, seeds from field plants were directly used for experiments after an after-ripening period of at least two months.

#### Phenotyping and growth conditions Experimental design

Unless described otherwise, experiments for phenotyping were conducted with twelve biological replicates per line. Four groups of three biological replicates were randomly assigned to one tray with 60 pots. The position of each group of three biological replicates within a tray was assigned randomly. In every phenotyping experiment, all trays were rotated 180° and moved every other day to minimize position effects. No block effects were detected on the measured traits (One-way ANOVA, Bonferroni corrected, p > 0.05).

#### Greenhouse growth conditions

Plants were grown under constant temperature of 22°C under long-day (LD) conditions [16 h light / 8 h dark]) and 65% humidity. Natural light was supplemented with LED arrays (Valoya, Model BX180c2, Spectrum AP67) to reach 120 - 150 µmol m^−2^ s^−1^ photosynthetic photon flux density. Seeds were stratified in water for five days at dark and 4°C. Around five to ten seeds were sown in each pot with ED73 potting mix (Einheitserdewerke, Sinntal-Altengronau, Germany). At the full expansion of the first two true leaves, plants were thinned to keep only one plant per pot. Because of reduced germination in some pots or severe damage of plants during thinning, the number of replicates per line varied.

#### Flowering time analysis

Flowering time was assessed through rosette leaf number (RLN), cauline leaf number (CLN) on the main shoot, and days to appearance of the floral bud (DTF). RLN and CLN were individually determined and combined to obtain total leaf number (TLN). DTF was consistently recorded, with a maximum one-day gap, representing the duration from germination to the emergence of the floral bud. Instances where flowering occurred later than 125 days or not at all were categorized as DTF 130, including the corresponding rosette leaf count at that time. Throughout the experiment, water status remained constant. In the experiment depicted in Fig. 1A and beyond, impaired germination hindered the analysis of one wildtype (8376), and RLN and CLN measurements were unattainable for seven additional wildtypes (eight in total), as the generative phase was not induced during our experiment. To incorporate these instances into the analysis, a DTF value of 130 was assigned.

#### Vernalization treatment for flowering time analysis

Seeds were allowed to germinate at 22°C under long days (LD, 16 hr light / 8 hr dark) and transferred to 4°C under short days (SD, 8 hr light / 16 hr dark) when cotyledons began to expand. After 60 days, the trays were transferred to the greenhouse with conditions described before. DTF values were reduced by 60 to account for the duration of the vernalization treatment.

#### Determination of plant growth

Plant growth was monitored daily by capturing top-view pictures using an EOS 2000D digital camera (Canon). Trays were identified by inclusion of triple-redundant QR codes. Images were normalized for size, orientation and perspective. This required between 15-45 s per image, where a quadrangle had to be placed at predetermined marker positions to compute a transformation matrix. A web-based tool was used for the interactive part (Labelbox, https://labelbox.com). Following normalization, a segmentation was performed, where the background was removed and the individual plants were extracted from the normalized multi-plant tray images. Background removal was performed by first applying a threshold on the images in the ‘Lab’ color space, followed by a series of morphological operations to remove noise and non-plant objects and a GrabCut-based postprocessing. The workflow was implemented in Python 3.6 and bash using OpenCV 3.1.0 and scikit-image 0.13.0 for image processing. Post-experiment, it was observed that the supplementary LED light in the greenhouse adversely affected image analysis. Consequently, images captured on days with the active supplemental lighting were excluded from the analysis. Moreover, plants with sizes below 5,000 pixels in the later stage of measurements and those smaller than 500 pixels on any day within the specified timeframe were entirely omitted from the dataset. These exclusions were made to eliminate likely empty pots or severely affected, dying plants (e.g., during thinning). Subsequently, area values in pixels underwent log transformation for subsequent analysis.

### Quantitative genetic analysis

#### Experimental design

F_2_ plants were cultivated in trays containing 60 pots, with four trays assigned to each F2 mapping population. Six plants from each parent line were randomly placed across the four trays. While all trays for a mapping population were kept together in the greenhouse, they underwent regular rotation, and the entire group of trays was relocated every second day to minimize positional effects. Growth and flowering time were analyzed following the procedures outlined in the “Flowering time analysis” section. Due to insufficient germination in certain pots, the F_2_ plant count per population varied.

#### DNA extraction

DNA was extracted according to (Yaffe et al. 2012) with Econospin® 96-well filter plates (Epoch Life Science, USA). The extraction buffer was modified to contain MES sodium salt instead of MES hydrate and RNase A (QIAGEN GmbH, Hilden, Germany).

#### WGS shallow sequencing

After DNA extraction and WGS library preparation, 150 bp paired-end reads were obtained on a HiSeq 3000 instrument (Illumina, San Diego, USA). The mean [±sd] number of reads of all F_2_ samples was 890,202 [356,734], ranging from 1,672 to 2,187,790.

#### Marker generation

To extract informative genetic markers, we enhanced the Trained Individual Genome Reconstruction (TIGER) CO analysis pipeline (Rowan et al. 2015), for compatibility with mapping populations of non-Col-0 (TAIR10) parents. The refined pipeline features an automated variant filtering step and a streamlined Snakemake-based process that outputs marker data compatible with the popular mapping package R/qtl (Broman et al. 2003; Rowan et al. 2015). The Snakemake pipeline, requiring minimal user input, generates cross-type marker input files suitable for the R library r/QTL (https://github.com/ibebio/tiger-pipeline).

Preprocessing involved trimming reads, mapping them to the TAIR10 reference, and removing duplicates. Variant calling for parent samples was performed with GATK, with the parent providing a higher number of variants selected as the source for alternative variant information (ALT). A “complete” variant file was obtained after soft filtering with GATK VariantFiltration (“QD < 5.0 || FS > 60.0 || MQ < 50.0 || MQRankSum < -12.5 || ReadPosRankSum < -8.0”) and extraction of biallelic SNPs. Subsequently, a “corrected” variant file was obtained through automated filtering, involving removal of variants deviating from the dominant peak of a bimodal Gaussian distribution fitted to QD values by SD*2.5, unimodal Gaussian distributions of FS by SD*2.5, of MQRankSum by SD*4 and MQ fixed at 50. Variants in centromeres, telomeres, and regions with transposable element annotations were excluded.

The complete marker file served as a reference for variant calling of F_2_ samples with GATK CollectAllelicCounts, excluding variants in organellar genomes and those with coverage deviating by five times the standard deviation of the mean. Samples with fewer than 7,000 variants were removed. Users can swiftly adjust these parameters through a configuration file. The prepared population-specific complete and corrected marker files, along with F_2_ allele count files, were employed as input for TIGER analysis, with slight script modifications to ensure file compatibility. The TIGER output files (one per F_2_ sample) were merged into a single R/qtl cross-type input file per population, and QC reports and genotype plots were generated at multiple steps during the pipeline.

#### QTL mapping

Phenotypic data was integrated into each R/qtl input file, and markers were filtered using r/QTL (Broman et al. 2003). F_2_ individuals with a low number of called markers, markers with substantial missing information, and individuals with very similar genotypes (90% similarity cutoff) were excluded. Markers with strong segregation distortion (P < 1e-7) and individuals with over 25 crossing-over events (more than twice than the expected median) were also removed. TIGER-identified crossing-over events served as unique markers in each F_2_ population, ensuring consistent genetic distances between markers. These prepared marker files were utilized for QTL mapping with a slightly modified version of the foxy QTL pipeline, which employs the R/qtl package in R (Feldman et al. 2017) (https://github.com/maxjfeldman/foxy_qtl_pipeline). The QTL mapping process involved several steps. A single QTL model genome scan utilized Haley-Knott regression to identify QTL with LOD scores exceeding the significant threshold, determined through 1,000 permutations at alpha = 0.05. A 2D genome scan employed a two-QTL model, with the significant threshold determined through 100 permutations at alpha = 0.05. A stepwise forward/backward selection procedure was then conducted to identify an additive, multiple QTL model based on maximizing the penalized LOD score (Broman et al. 2003; Feldman et al. 2017). Following the concatenation of all QTL tables for all populations and traits, physical positions were assigned to each QTL, along with their respective 95% Bayes intervals.

### qRT expression analysis

As described in Ruffley et al. 2024. All primer sequences are listed in Table S8.

### RNA-seq

Normalized RNA from the qRT experiment was used for mRNA library prep using an in-house custom protocol adapted from Illumina’s TruSeq library prep, with details provided in (Cambiagno et al. 2021). FASTQ files from multiple lanes were merged and mapped to the TAIR10 transcriptome using RSEM (bowtie2, version 2.2.3) with default parameters. A mRNA counts file was obtained with featurecounts (version 1.6.1). One of three biological replicates of line ID9402-01-R05 (replicate 3) was removed due to an insufficient number of feature counts (690569). Differential expression analysis was conducted with edgeR, with model.matrix(∼0+group) as design and the function makeContrasts and glmTreat to retrieve mutant-wildtype contrast specific lists of DEGs (FDR<0.1 and |log2 FoldChange|>1). GO enrichment analysis was conducted with ShinyGO with the pathway database “GO Biological Process” with 22157 gene IDs as background and a FDR cutoff of 0.01 (Ge et al. 2020).

### *FT* and *FLC* sequence analysis

SNP variants at *FT* as available from the 1001 Genomes resource (http://1001genomes.org/data/GMI-MPI/releases/v3.1/) were extracted using VCFtools (version 0.1.16) with “--chr 1 --from-bp 24325373 --to-bp 24335992”. VCF files were transformed to fasta using PGDSpider (version 2.1.1.5) and aligned with muscle (version 3.8.31). A neighbor-joining tree was built with MEGA X (Kumar et al. 2018). The dataset comprised 1,135 sequences. All positions with fewer than 95% coverage were eliminated, resulting in 382 positions. A tree was visualized with iTOL (Letunic and Bork 2021). Variants at *FLC* were extracted using VCFtools “--chr 5 --from-bp 3173382 --to-bp 3179448” and filtered for SNPs with a minor allele frequency of 10% and maximum missing data of 10% with “--remove-indels –maf 0.1 --max-missing 0.9”, resulting in 35 variants. Missing genotypes were imputed with Beagle 4.0 (version r1399) and a Minimum Spanning Network was constructed with PopArt (Bandelt et al. 1999). The *FLC* expression data were from (Kawakatsu et al. 2016) and the flowering time data from https://arapheno.1001genomes.org/phenotype/262/.

### Measurement of nitrogen isotope ratio and carbon and nitrogen content

Mutants and wild types were grown in three pots representing three biological replicates grown at 22°C under LD in the greenhouse (similar conditions described in section “Flowering time analysis”). The pots were randomly distributed over a total of ten trays, which were rotated and moved every day to reduce position effects. After allowing germination and establishment of the first true leaves, plants were thinned to three plants per pot. The leaves of different plants never overlapped with each other during the experiment. Rosettes were harvested at the initiation of flowering or after 22 days after germination (whatever was first). Depending on the rosette size at the initiation of flowering several plants were combined to one replicate to reach the required amount of tissue for analysis. Plant tissue was dried at 60°C for 24 hours and homogenized in 5 ml tubes (Eppendorf, Germany) containing 5 ball bearings to very fine and uniform powder. Dried material was transferred to a 1.5 ml microfuge tube and send to Isolab GmbH (Schweitenkirchen, Germany) for an analysis of nitrogen isotope composition and C and N content (δ^15^N, %N and %C) with ^15^N-CF-IRMS and ^13^C-CF-IRMS. Four technical replicates per sample were analyzed. For a more detailed description of the procedure, see (Werner and Roßmann 2015). Data are presented as δ^15^N [‰] vs. AIR, and percent mass carbon (%C) and nitrogen (%N) in the plant tissue, respectively.

## Acknowledgements

We thank the Weigel Lab members for comments and discussion. We thank S. Schäfer, N. Betz, A. Duque, A. Spazierer, F. Strauss, N. Vasilenko, J. Kreienbrink, and A. Ziewanah for technical assistance.

## Author Contributions

Conceptualization - UL; Methodology – UL, IB, RS, WY; Experiment – UL, RS, WY, MK; Data analysis

– UL; Writing – original draft: UL; Writing – review & editing – all authors.

## Data and Material Availability

Reads were deposited at NCBI SRA with accession number: <TBD upon publication>. Seeds were deposited at NASC with the accession number <TBD upon publication>.

## Code Availability

The TIGER snakemake pipeline is available at https://github.com/ibebio/tiger-pipeline. All tools used for the bioinformatic analyses are publicly available. Unless specified otherwise, default parameters were used.

