## Supplemental Figures for "Cryptic Variation in Adaptive Phenotypes Revealed by Panspecific *flc* Mutants"

### Supp. figure 1

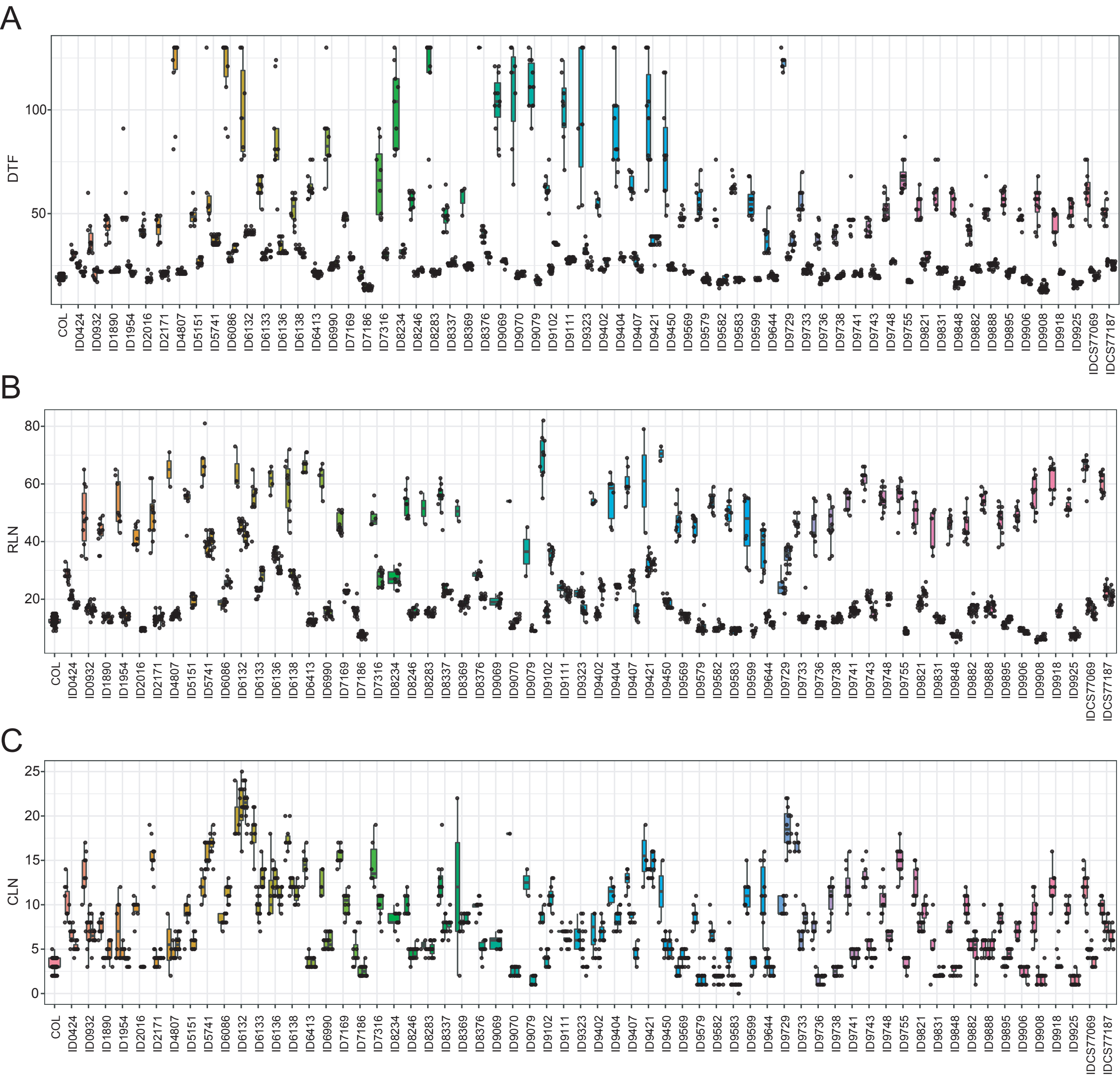

### Supp. figure 2

A

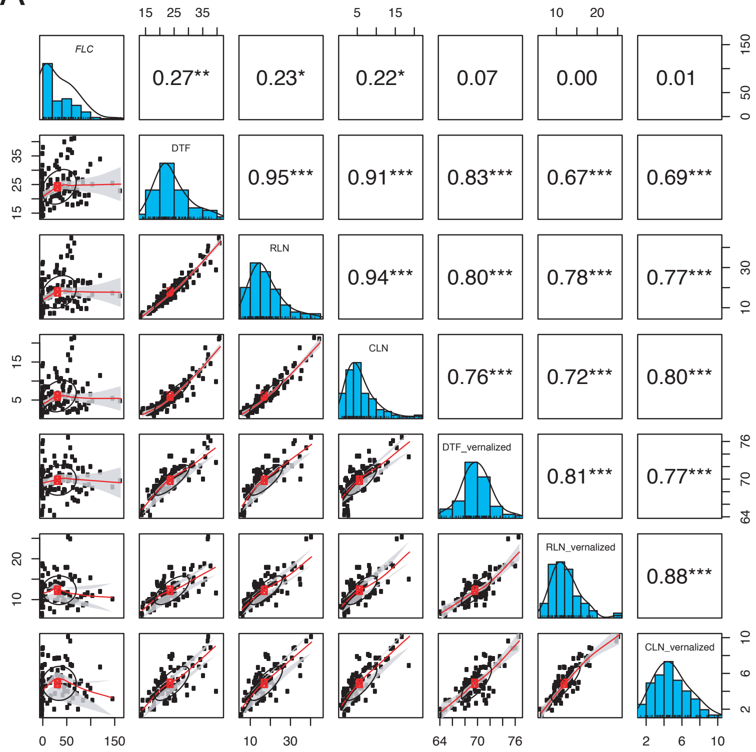


B

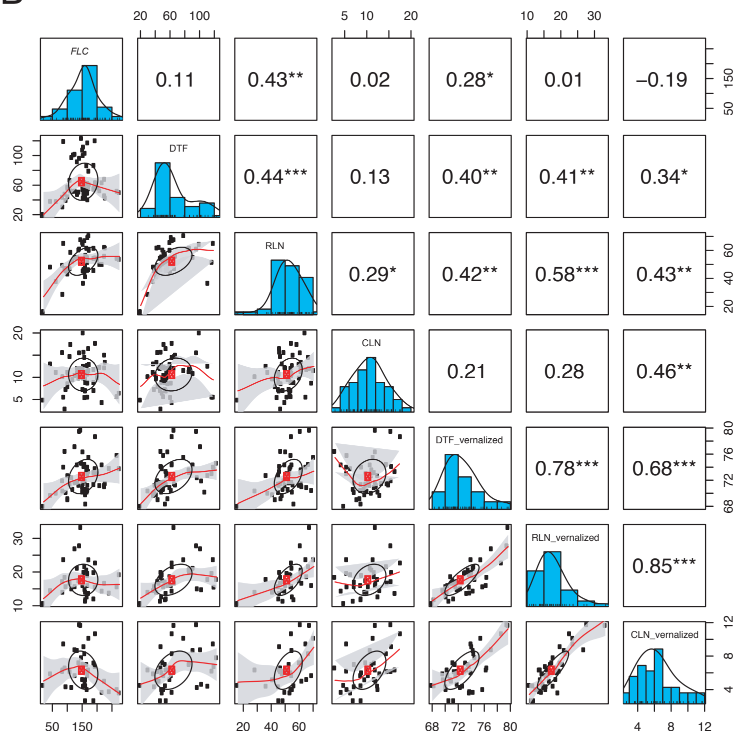

Supp. figure 3

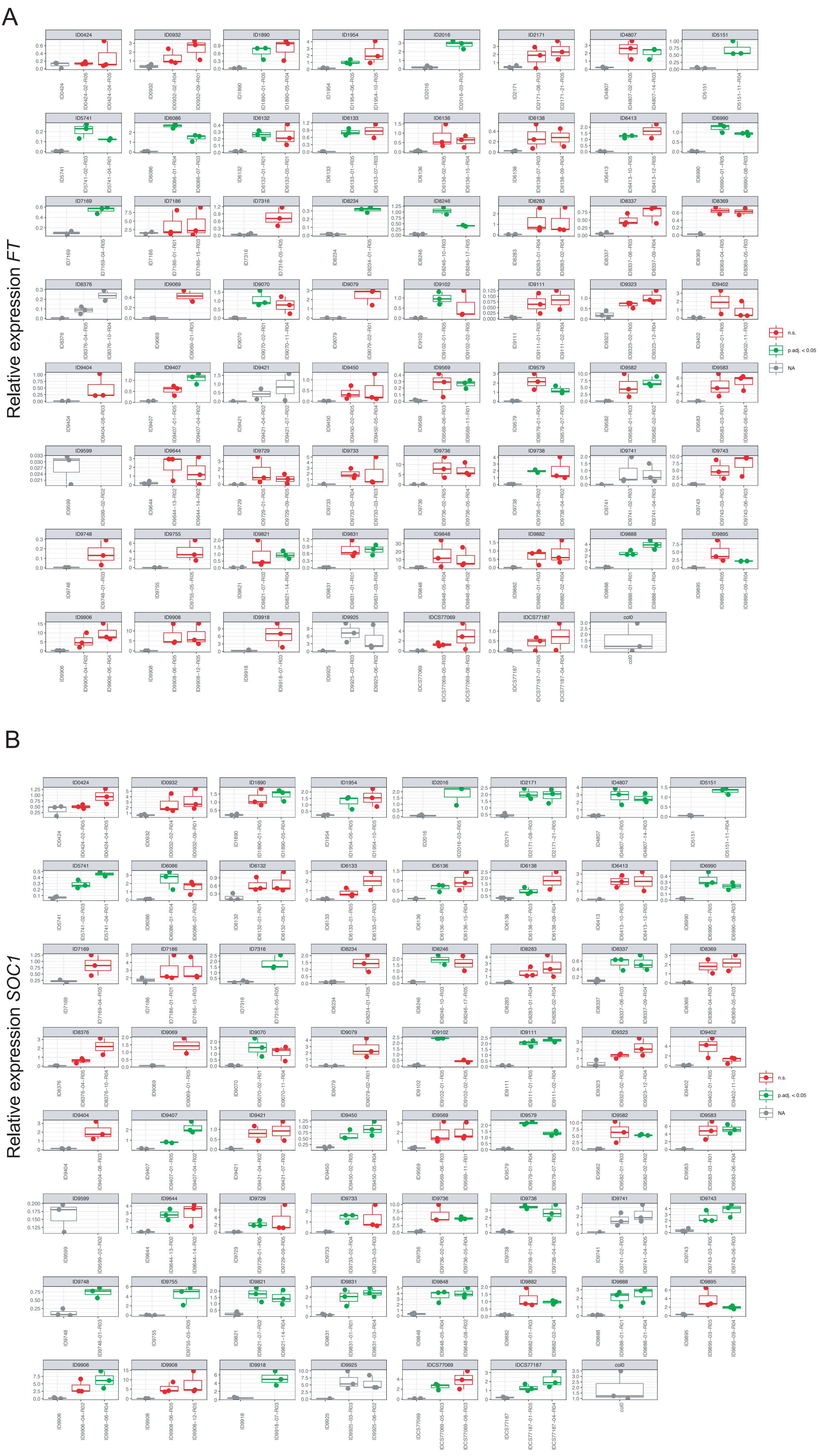

Supp. figure 4

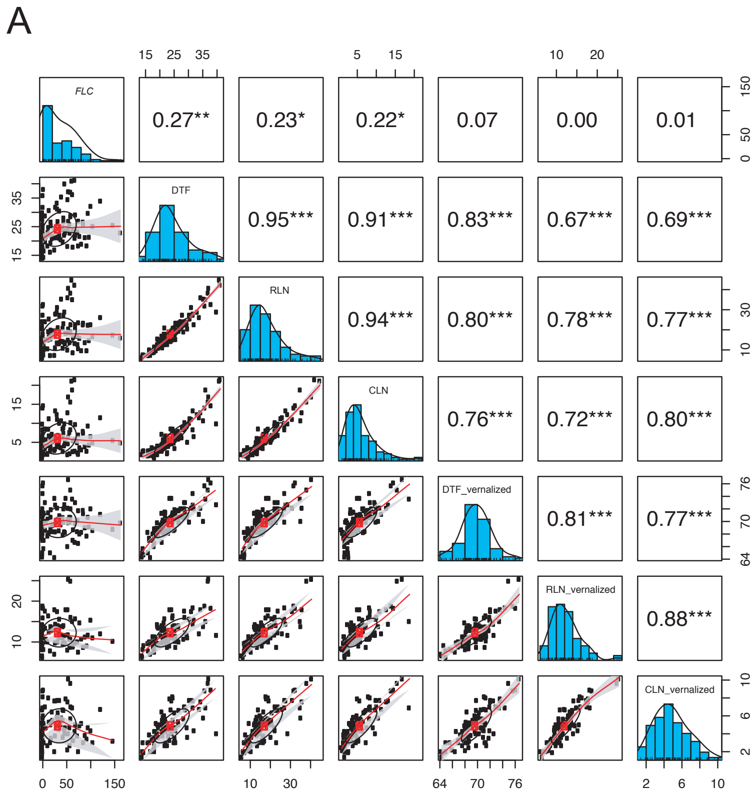


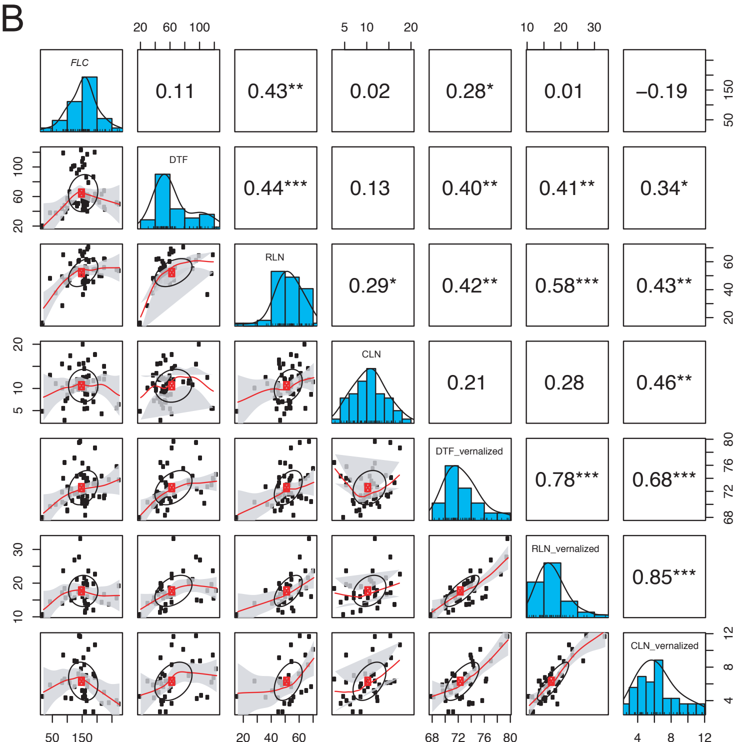

### Supp. figure 5

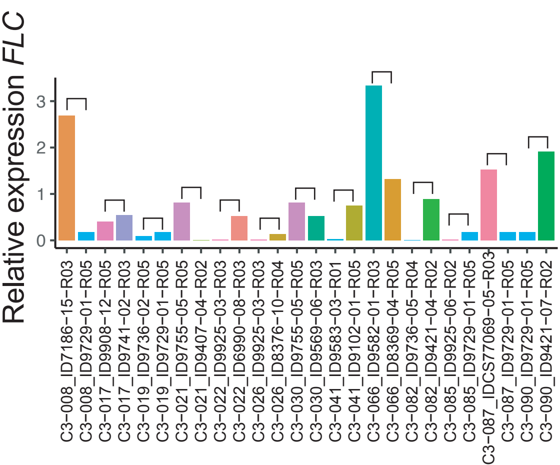

Supp. figure 6

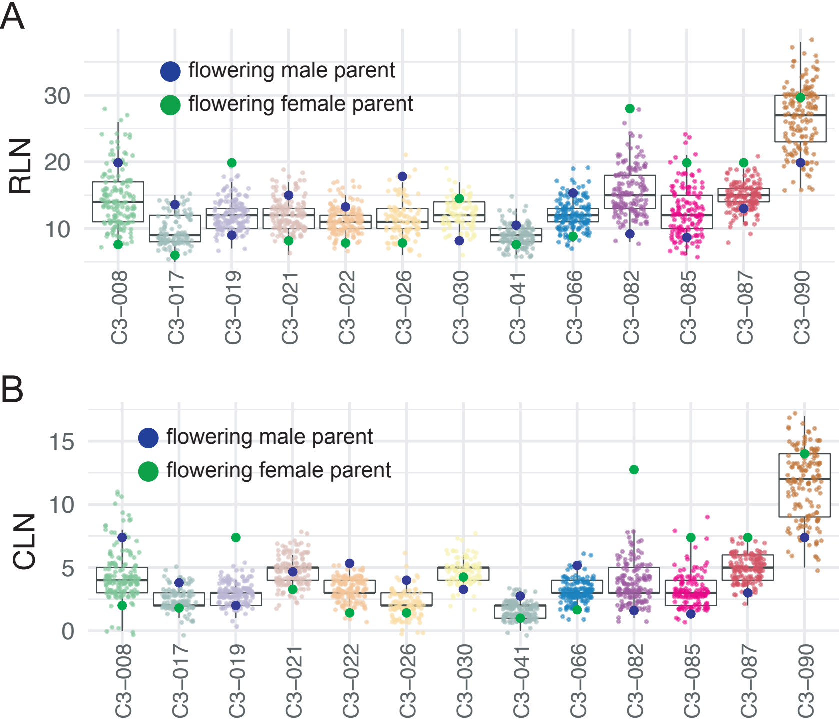

### Supp. figure 7

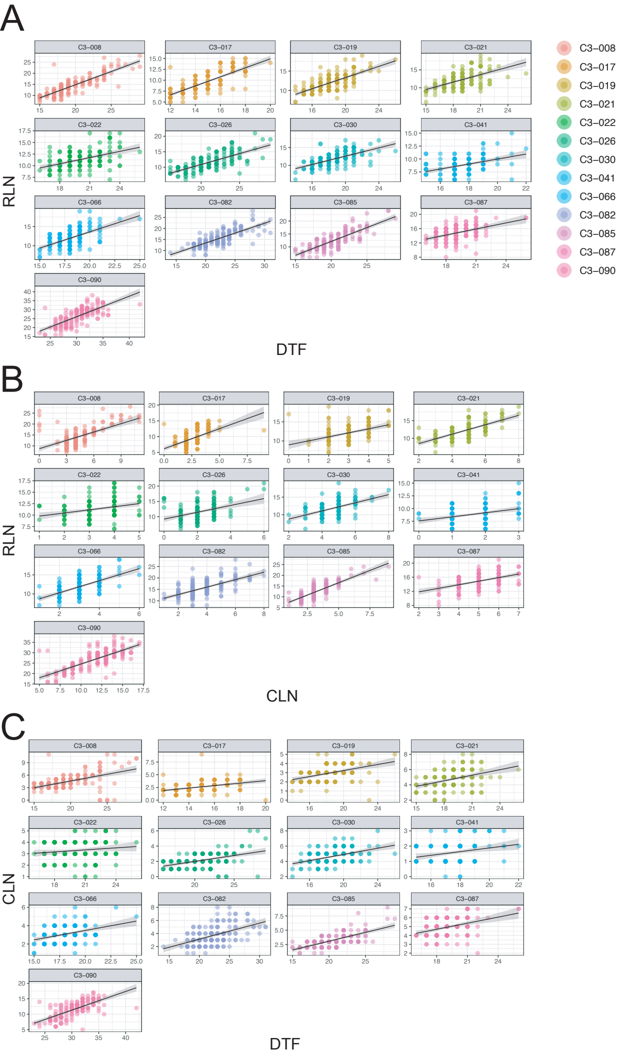

### Supp. figure 8

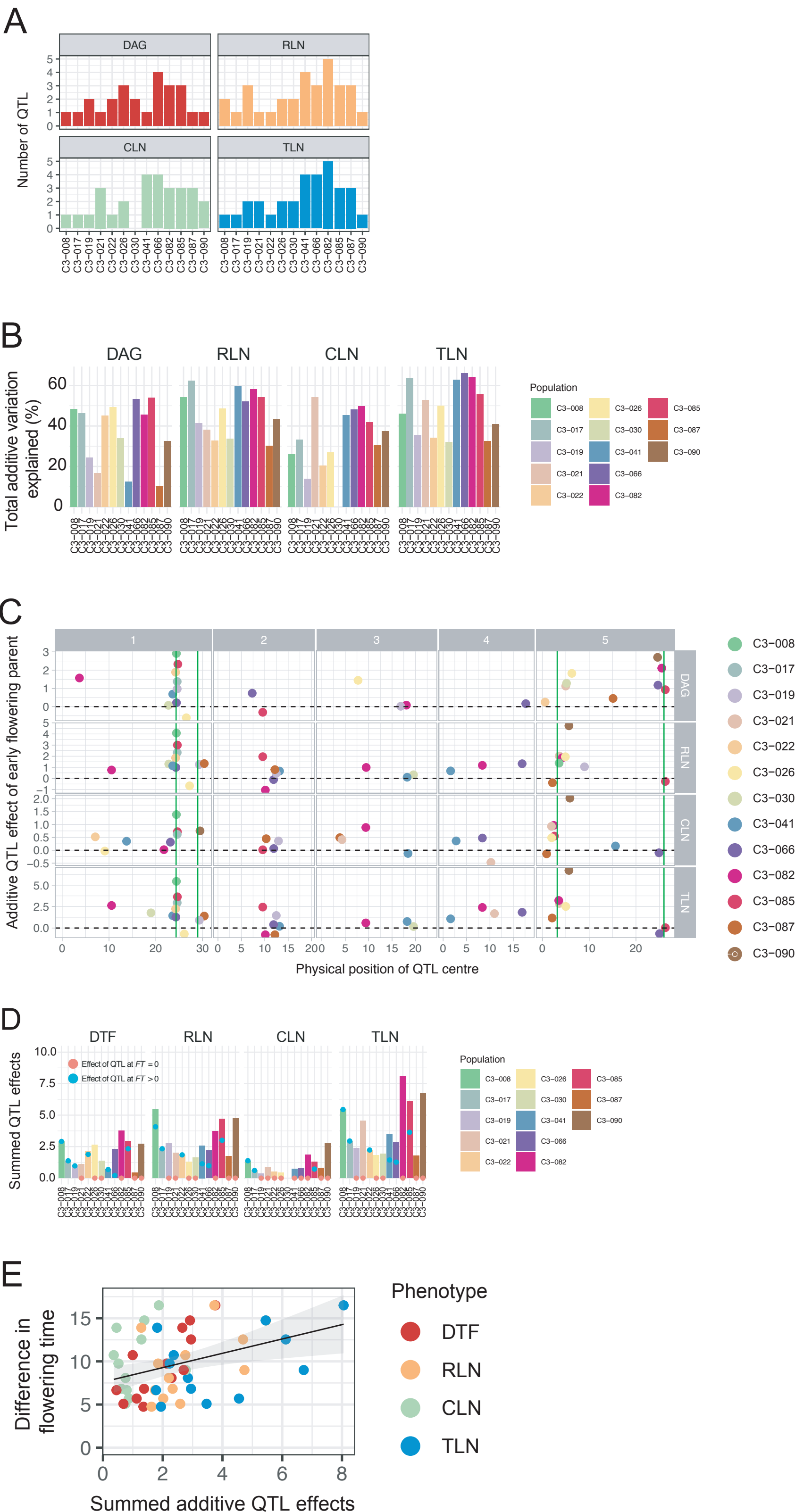

### Supp. figure 9

A

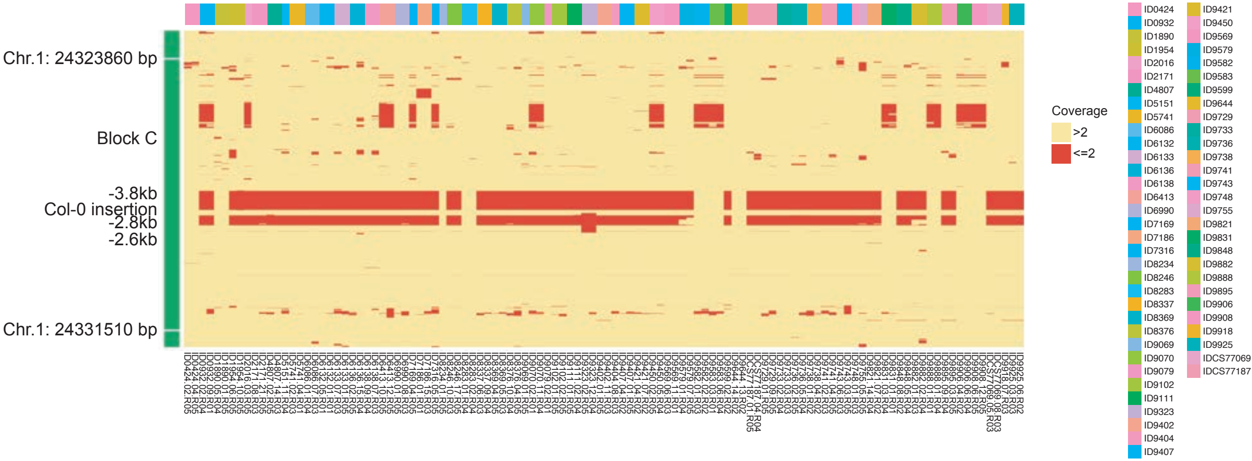


B

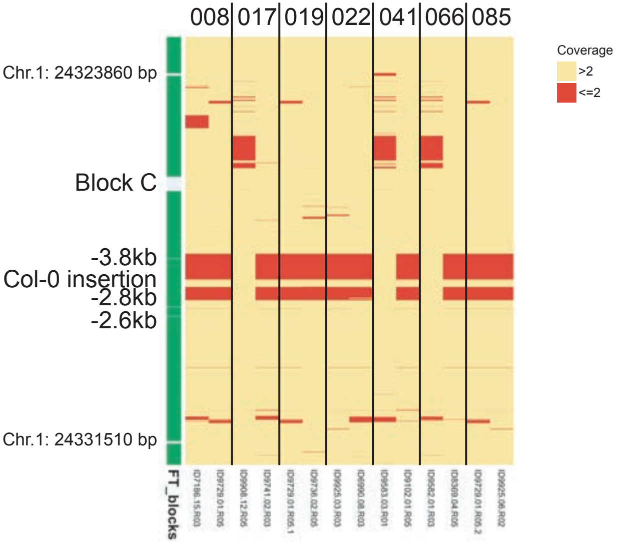

Supp. figure 11

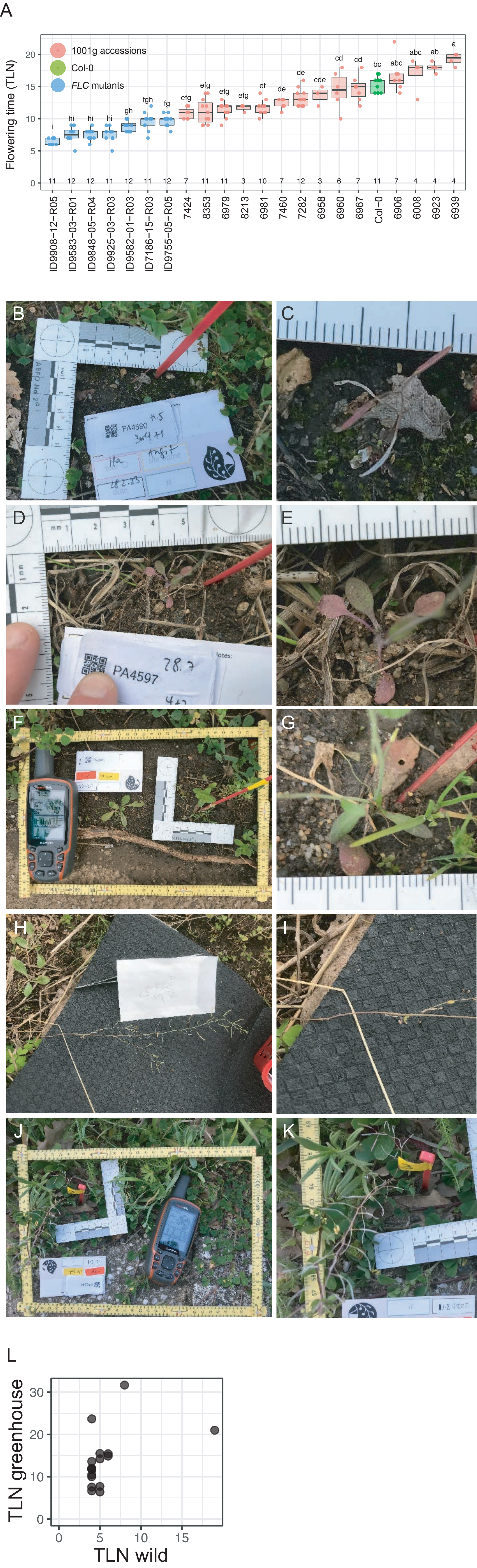

Supp. figure 11

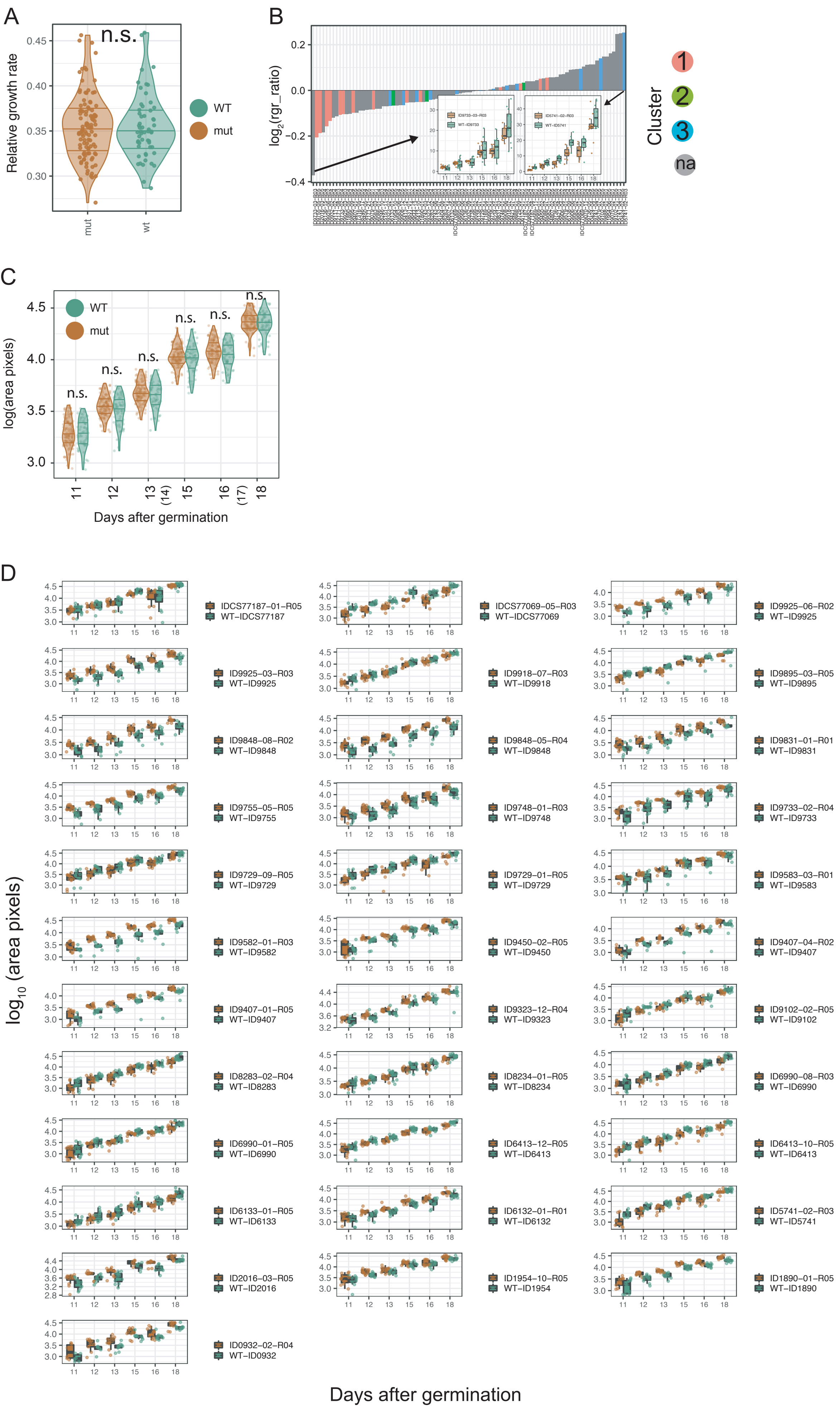

### Supp. figure 12

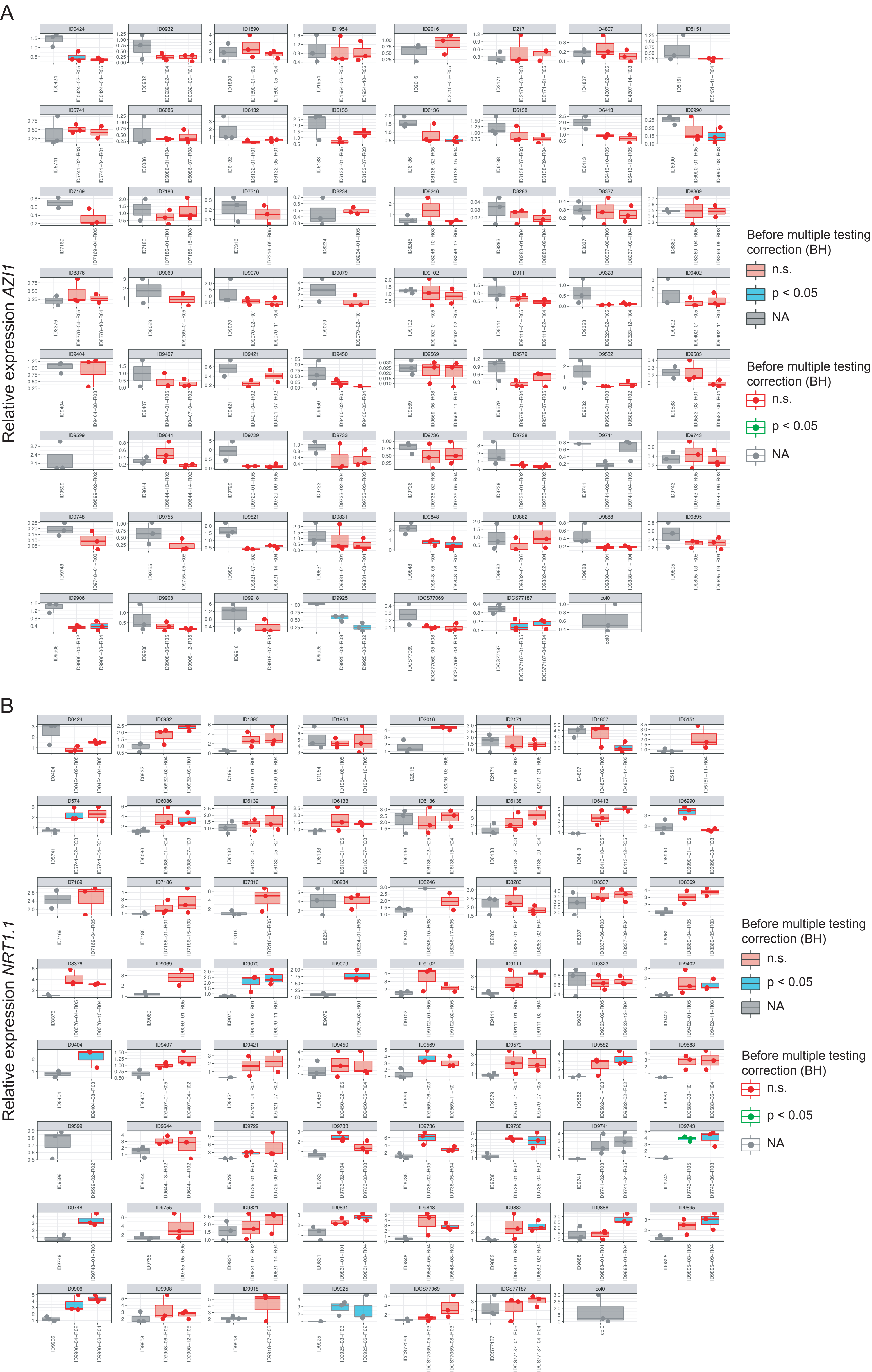

### Supp. figure 13

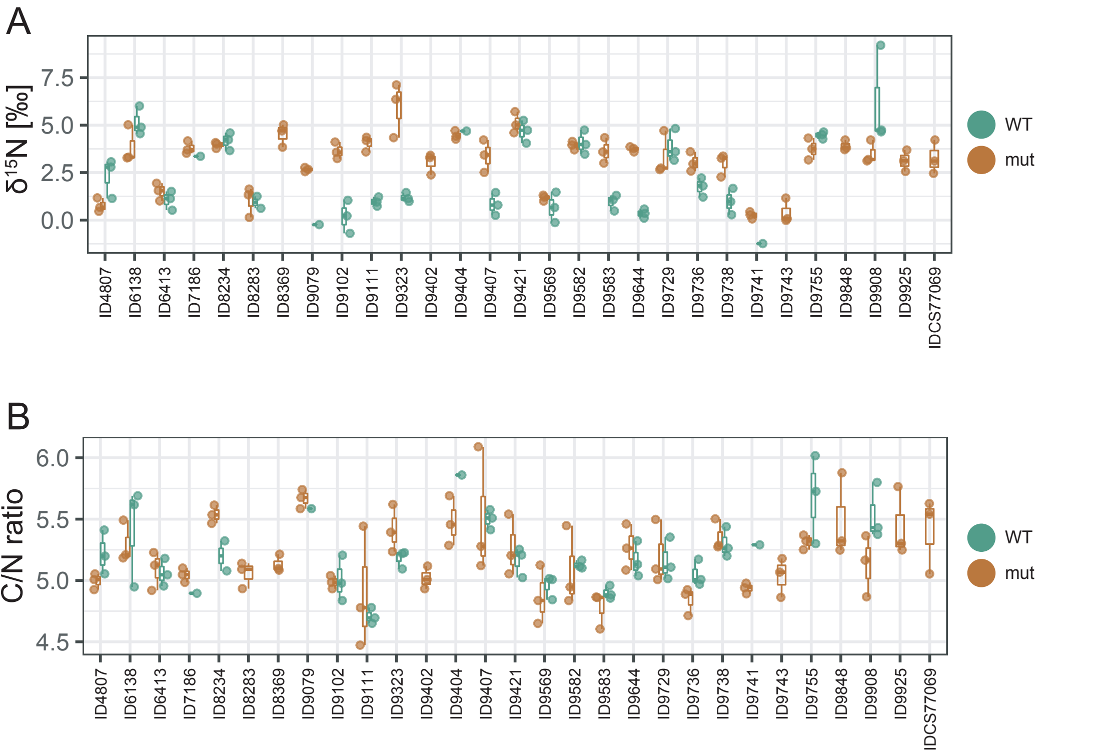
